# Single-cell analysis of megakaryopoiesis in peripheral CD34^+^ cells: insights into ETV6-related thrombocytopenia

**DOI:** 10.1101/2022.09.20.508634

**Authors:** Timothée Bigot, Elisa Gabinaud, Laurent Hannouche, Véronique Sbarra, Elisa Andersen, Delphine Bastelica, Céline Falaise, Manal Ibrahim-Kosta, Marie Loosveld, Paul Saultier, Dominique Payet-Bornet, Marie-Christine Alessi, Delphine Potier, Marjorie Poggi

## Abstract

Expansion of human megakaryoblasts from peripheral blood-derived CD34^+^ cells is commonly used to characterize inherited or acquired thrombocytopenia and evaluate defects in megakaryocyte (MK) differentiation, MK maturation and proplatelet formation. We applied single-cell RNA sequencing to understand local gene expression changes during megakaryopoiesis (days 6 and 11 of differentiation) in peripheral CD34^+^ cells from healthy controls and patients with *ETV6*-related thrombocytopenia.

Analysis of gene expression and regulon activity revealed distinct clusters partitioned into seven major cell stages: hematopoietic stem/progenitor cells (HSPC), common-myeloid progenitors (CMP), MK-primed CMP, granulocyte-monocyte progenitors, megakaryocyte-erythroid progenitors (MEP), MK progenitor /mature MK (MKP/MK) and platelets. We observed a subpopulation of MEP that arose directly from HSPC, deviating from the canonical MK differentiation pathway.

ETV6 deficiency was characterized by an increase in HSPC, a decrease in MKP/MK, and a lack of platelets. ETV6 deficiency also led to the development of aberrant MEP and MKP/MK cell populations. Genes involved in “mitochondrial” and “DNA repair” pathways were downregulated, while genes involved in “translation” pathways were upregulated. Analysis of patient samples and hematopoietic cell lines transduced with an *ETV6* variant revealed increased translation in MK. Ribosomal protein small 6 (RPS6) levels in MK, platelets and peripheral blood mononuclear cells was consistent with the translation findings.

Our results provide a framework to understand peripheral CD34^+^ cell-derived megakaryocytic cultures. Our observations also shed light on *ETV6*-variant pathology and reveal potential targets for diagnostic and therapeutic purposes.

**Key points:**

- scRNAseq gain insight into *in vitro* megakaryopoiesis, identify MK-primed CMP, and a differentiation trajectory that bypasses the CMP.
- *ETV6* variants led to the development of aberrant MEP and MK cell populations.

## Introduction

Megakaryocytes (MK) are fragile and only represent a small fraction of normal bone marrow cells (approximately 0.05% of mononuclear cells), which has hindered the study of megakaryopoiesis and the hierarchical structure of these cells. Most studies have analyzed flow-sorted cell populations, which limits assessments to a predefined cell subset.

*In vitro* culture systems for MK progenitors has enabled the analysis of megakaryopoiesis and regulation of MK differentiation. *In vitro* platelet production may represent an alternative to at least partially compensate for the increasing demand for platelet concentrates. Many investigators have attempted to increase platelet production *in vitro* by modifying culture media components^1–4^. An *ex-vivo* serum-free liquid culture system has also been used to expand normal human megakaryoblasts from purified peripheral-derived CD34^+^ cells to characterize acquired or inherited thrombocytopenia (IT) and evaluate defects in MK differentiation, maturation and proplatelet formation^5,6^. Furthermore, these models are largely used to evaluate the effect of infectious diseases or therapeutic agents on megakaryoblast differentiation^7–10^ and investigate novel mechanisms in MK differentiation and platelet function^11–16^

In the current study, we aimed to characterize the developmental stages of normal *in vitro* megakaryopoiesis. Primary CD34^+^-hematopoietic progenitor cells were induced to differentiate along the megakaryocytic lineage in liquid suspension cultures continuously exposed to thrombopoietin. We compared our results to those obtained in patients with ETV6-related thrombocytopenia (ETV6-RT), a highly penetrant form of IT with autosomal dominant inheritance^17^. The common phenotype observed in ETV6-RT includes moderate thrombocytopenia sometimes associated with bleeding and predisposition to acute T or B-cell lymphoblastic leukemia. ETV6-RT is also associated with B-cell lymphoma, acute myeloid leukemia and myelodysplasia to a lesser extent^18–20^.

The precise role that *ETV6* plays in megakaryocyte differentiation remains poorly understood. Studying *Etv6* function in murine models is challenging because complete loss of the gene is lethal^21,22^ and heterozygous *Etv6* mice have obvious unperturbed hematopoiesis^23^. Human induced pluripotent stem cells (iPSC) harboring a pathogenic heterozygous *ETV6* mutation do not give rise to an increase in hematopoietic progenitor cells and MK. However, iPSC carrying the homozygous *ETV6* mutation give rise to an increase in hematopoietic progenitor cells and immature MK, as observed in heterozygous patients using an *in vitro* model of CD34^+^-derived MK^24,25^. Therefore, further investigation using patient cells is required to better understand defective ETV6-megakaryopoiesis.

Using cells derived from controls and patients harboring the *ETV6*-variants, we applied single-cell RNA sequencing to examine the transcriptome of each cell type during differentiation from hematopoietic stem/progenitor cells (HSPC) to MK. We observed classical hematopoietic cell populations (HSPC, common-myeloid progenitors (CMP), granulocyte-monocyte progenitors (GMP), megakaryocyte-erythroid progenitors (MEP), MK progenitors (MKP), mature MK and platelets (Plt) and an unusual MK-primed CMP subpopulation. We observed a subpopulation of MEP that arose directly from HSPC, bypassing CMP. ETV6 deficiency led to the development of aberrant MEP and MK populations, with an enrichment in “mitochondrial”, “DNA repair” and “translation” pathways. Our findings provide insight into peripheral CD34^+^-megakaryopoiesis, *ETV6*-variant pathology and potential targets for diagnostic and therapeutic purposes.

## Methods

### *In vitro* megakaryocyte differentiation

ETV6-variant carriers and healthy volunteers were recruited at the Center for the Investigation of Hemorrhagic and Thrombotic Pathologies at Marseille University Hospital (authorization number 20200T2-02). Peripheral CD34^+^-cells were purified using magnetic cell sorting (Miltenyi-Biotec) and then cultured in StemSpan Serum-Free Expansion Medium combined with Megakaryocyte Expansion Supplement (Stemcell Technologies)^26^.

### Single-cell RNA sequencing

Control (n=2) and ETV6-defective (n=2) cells were harvested from culture at days 6 and 11 (Figure 1A). The cell samples were labeled with a distinct hashtag oligo (TotalSeq, Biolegend) and pooled. Single-cell isolation was then carried out with the 10X Genomics Technology using the Chromium Next GEM Single Cell 5’ Kit v2. Single-cell cDNA synthesis and sequencing libraries were prepared with single-cell 5’ Library and Gel Bead kit.

**Figure 1.**
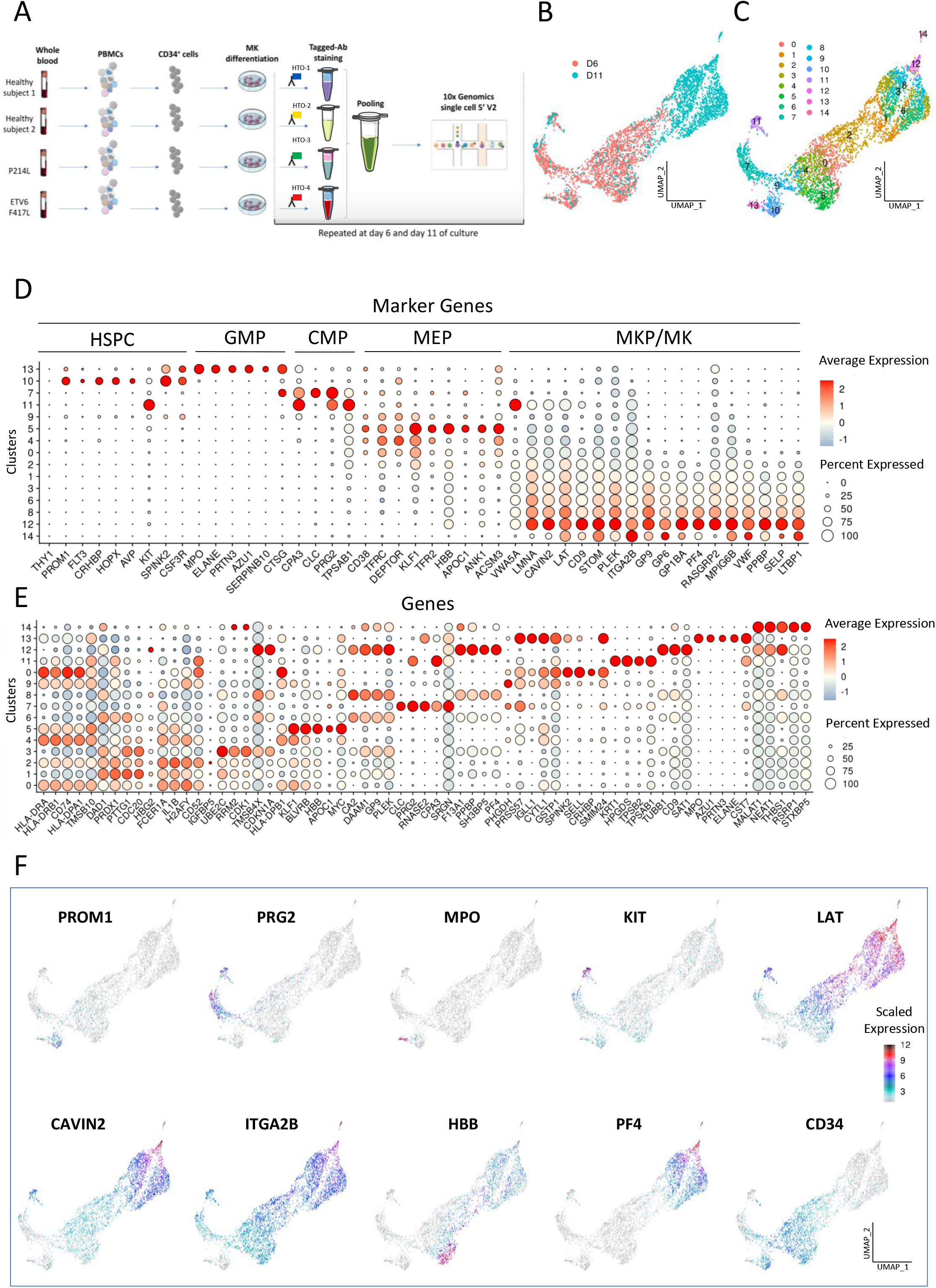
Single-cell RNA sequencing analysis in megakaryopoiesis cell stages in CD34^+^ cells isolated from healthy volunteers. **A-** Schematic diagram of the experimental design. Fresh CD34^+^ cells from human peripheral blood were isolated via density gradient and magnetic cell sorting (n=2 healthy volunteers). CD34^+^ cells were differentiated into megakaryocytes using serum-free StemSpan SFEM II medium with megakaryocyte expansion supplement (SCF, IL6, TPO and IL9) (STEMCELL Technologies). Cells from different samples are incubated with DNA-barcoded antibodies recognizing ubiquitous cell surface proteins. Distinct barcodes (referred to as hashtag-oligos, HTO) on the antibodies allow pooling of multiple samples into one single-cell RNA (scRNA) sequencing experiment. The cells were analyzed at differentiation days 6 and 11 using 10X genomics.. **B-** UMAP plot of the control cells on days 6 and 11 after merging the two data sets (D6 and D11, respectively). The cells analyzed on day 6 are shown in orange, and the cells analyzed on day 11 are shown in blue. **C-** UMAP plot of the control cells on days 6 and 11 (D6 and D11, respectively), with cell clustering performed using a resolution of 1.2. Each cluster is indicated with a distinct color. **D-** Seurat dot plot showing the expression levels of known specific hematopoietic cell marker genes in each cluster. **E-** Seurat dot plot of the top five differentially expressed genes in each cluster. Dot size represents the percentage of cells expressing the gene of interest (percent expressed), while the color gradient represents the scaled average expression of the genes in each cluster (a negative value corresponds to expression levels below the mean). **F-** Feature plot showing the expression of genes representative of each cell type (*PROM1, PRG2, MPO, KIT, LAT, CAVIN2, ITGA2B, HBB, PF4* and *CD34*).

### Additional methods

See the supplemental methods for additional details regarding data preprocessing and analysis, site-directed mutagenesis, western-blot, microscopy, flow-cytometry, transduction and statistical analyses.

## Results

### Characterization of differentiation in control CD34^+^-cells

We performed UMAP (Uniform Manifold Approximation and Projection) non-linear dimensional reduction to analyze cell transcriptome heterogeneity. The observed variation between the transcription profiles of cells after 6 and 11 days in culture indicated that distinct gene sets were involved in each stage of differentiation (Figure 1B).

Using unsupervised clustering, we identified a total of 15 clusters (Figure 1C), which were present in both controls (Supplemental figure 1A-C). We performed a detailed characterization of each cell cluster based on known cell gene sets (signature)^27–30^ (Figure 1D) and the top differentially expressed genes (DEGs) (Figure 1E; Supplemental Figure 2, Supplemental Table 1).

Cluster 10 was designated HSPC based on the expression profiles of *PROM1, CRHBP, FLT3, HOPX* and *AVP*. Similarly, clusters 11 and 7 were designated CMP based on the expression profiles of *CPA3, PRG2, CLC* and *TPSAB1*. Cluster 11 differed from cluster 7 as it shared marker genes with the MKP/MK clusters (*VWA5A, LMNA, CAVIN2, LAT, CD9* and *ITGA2B*) (Figures 1D-F) and expressed the genes *KIT, KRT1, HPGDS* and *TPSB2*. We thus defined this cluster 11 as MK-primed CMP. Cluster 13 was designated GMP because it expressed specific markers such as *MPO, ELANE* and *PRTN3*. Clusters 9, 5, 4, 0 and 2 were designated MEP. Although these clusters shared a strong common MEP signature (*CD38, TFRC, DEPTOR, KLF1, TFR2*), we further characterized these five subpopulations. Cluster 9 was the most immature population and was designated “early MEP” based on the remaining expression levels of HSPC and CMP genes. Cluster 5 was designated a MEP-ERP subpopulation based on the genes associated with erythroid progenitors (ERP) such as *HBB, TFR2* and *ANK1*. Clusters 4, 0 and 2 were designated a MEP-MK subpopulation based on the expression levels of MK genes, such as *GP9, GP6, GP1BA* and *MPIGB*, and the gradual increase in *MYH9* expression levels between clusters 4, 0 and 2 (Figures 1E-F, Supplemental figures 1D and 2). Clusters 1, 6, 3, 8 and 12 were primarily observed at day 11 and were designated MK based on the expression of genes associated with classical MK. Specific MK markers (*GP6, GP9, PF4, P2RY1*) gradually increased from cluster 1 to 12 (Figure 1D, Supplemental figures 1D and 2). Of these five clusters, cluster 1 was the most immature stage (MKP) and cluster 12 was the most mature. Cluster 14 was only observed at day 11 was designated platelets based on the low number of genes expressed as compared with the MKP/MK populations (mean±SD: 1,418±347 vs. 4,208 ±1,399) and overall RNA count (mean±SD: 2,243±751 vs. 23,855±14,826) (Figures 2A-B, Supplemental figure 1D). Furthermore, cluster 14 expressed the same genes as cluster 12, with higher expression levels of *GP6* and *ITGA2B* (Figures 1D-E, Supplemental figures 1D and 2).

**Figure 2.**
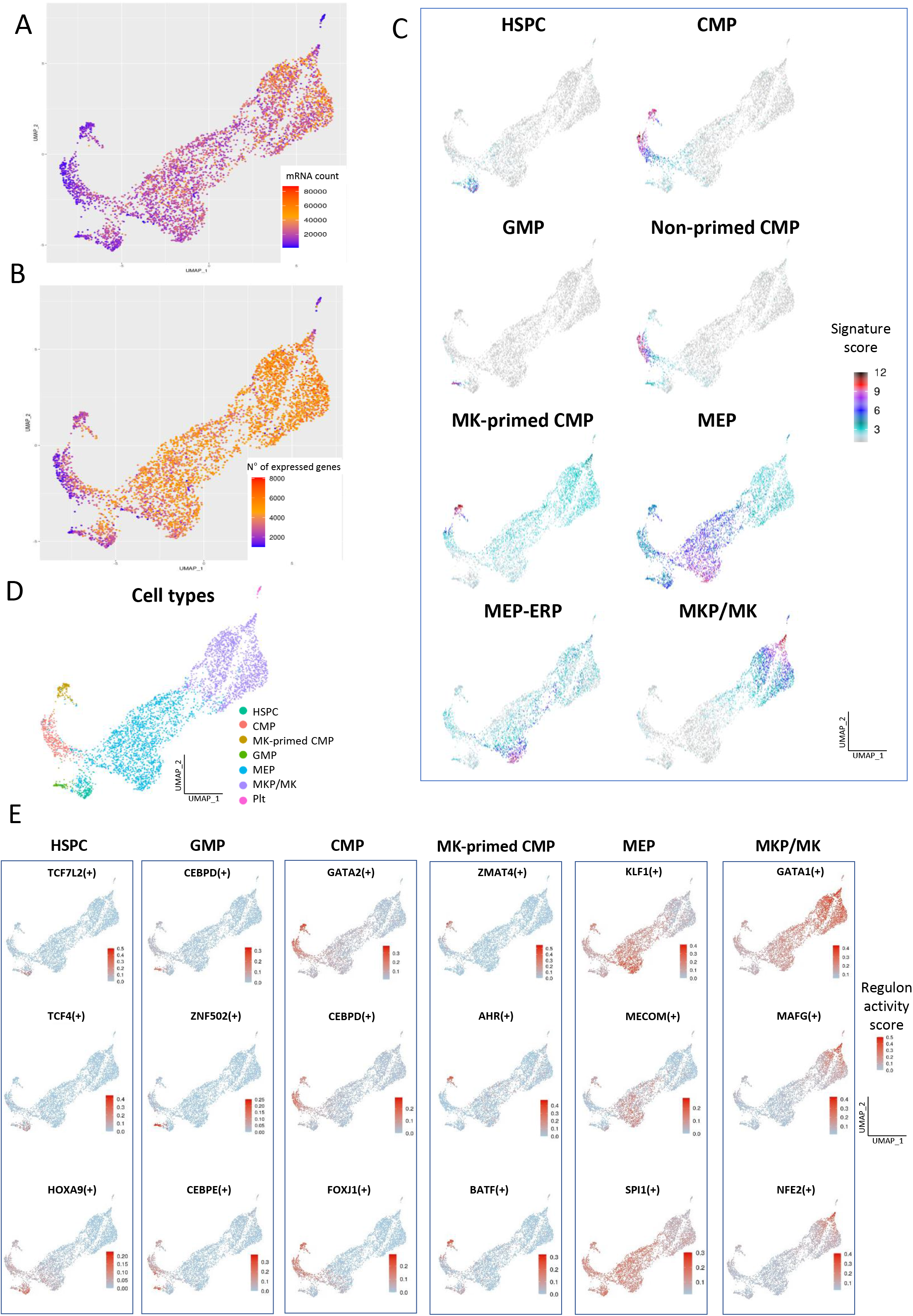
CD34^+^ cell-derived cultures contain cells of various hematopoietic states. A- UMAP plot showing RNA count for hematopoietic cells derived from normal CD34^+^ cells. B- UMAP plot showing the number of genes expressed for hematopoietic cells derived from normal CD34^+^ cells. C- Feature plots showing the expression of lineage signature gene set score for hematopoietic cells derived from normal CD34^+^ cells. D- UMAP plot of hematopoietic cells derived from normal CD34^+^ cells color-coded according to cell type. Numbers correspond to clusters. E- UMAP plots displaying Top 3 regulon activity enriched in each lineage: HSPC (TCF7L2, TCF4 and HOXA9), GMP (CEBPD, ZNF502 and CEBPE), CMP (GATA2, CEBPD and FOXJ1), MK-primed CMP (ZMAT4, AHR and BATF), MEP (KLF1, MECOM and SPI1) and MKP/MK (GATA1, MAFG and NFE2).

For each cell type, we proposed enriched gene signatures that included known cell-type specific genes, top-ranked DEGs and selective genes primarily expressed in cell stage-related clusters (Supplemental table 2). Lineage signature scores were then computed and assigned to each cell (Figure 2C). Cell types were then defined accordingly (Figure 2D).

### Inference of megakaryopoiesis regulon activity in control cells

The cell state transitions in megakaryopoiesis are tightly controlled by transcription factors (TF)^17^. Regulons are inferred groups of genes controlled as a unit by the same repressor or activator TF^31^. For each regulon, the activity score was calculated based on the cellular expression values for all genes. Cell type-specific regulons provide an opportunity to identify key regulators of cell fate decisions and establish cell signatures. Analysis of our control dataset using the single-cell regulatory network inference and clustering (SCENIC) workflow provided insight into cell type-specific regulons that drive cellular heterogeneity. Comparing each cell-types, we analyzed the top regulons to isolate key regulons at each stage of differentiation (Figure 2E, Supplemental figure 4). All 312 identified regulons are available in Supplemental table 3.

### Characterization of the *ETV6* variants

We performed a functional study of the novel *ETV6* variant pF417LTer4 compared with the previously described p.P214L^25^ (Supplemental figure 5A). The nonsense mutation p.F417LTer4 is located in the ETS domain, while the missense mutation p.P214L is localized in the linker domain (Supplemental figure 5B). The clinical and laboratory characteristics of the patients are shown in Supplemental table 4.

We analyzed the repressive activity of the two variants using a dual-luciferase reporter assay. Co-transfection of the reporter plasmid containing the ETS-binding site along with expression of a plasmid encoding wild type (WT) *ETV6* resulted in almost 90% inhibition of luciferase activity. Substitution of WT *ETV6* with any of the *ETV6* variants led to a significant reduction in repressive activity (85% to 100%) (Supplemental figure 5C). Western-blot analysis showed that mutant ETV6 protein was expressed in the GripTite293 macrophage scavenger receptor (MSR) cell line (Supplemental figure 5D).

Subcellular fractionation of GripTite293 MSR cells showed increased ETV6 protein levels in the cytoplasmic fraction and decreased levels or absence of ETV6 in the nuclear fraction in cells expressing the p.P1214L or p.F417Lter4 variants compared with cells expressing the WT protein (Supplemental figure 5E). Microscopy confirmed that WT ETV6 concentrated primarily in cell nuclei, whereas both ETV6 variants were predominantly localized in the cytoplasm (Supplemental figure 5F).

### Single-cell transcriptional profiling of *ETV6*-variant CD34^+^-cells during megakaryopoiesis

We performed UMAP non-linear dimensional reduction to analyze cell transcriptome heterogeneity and clustering. Compared with control cells, the day 6 and day 11 transcriptome profiles of patient cells overlapped much more, thereby suggesting that differentiation was delayed with accumulation of early-stage cells in both *ETV6*-variant carriers (Figure 3A, Supplemental figure 6A).

**Figure 3.**
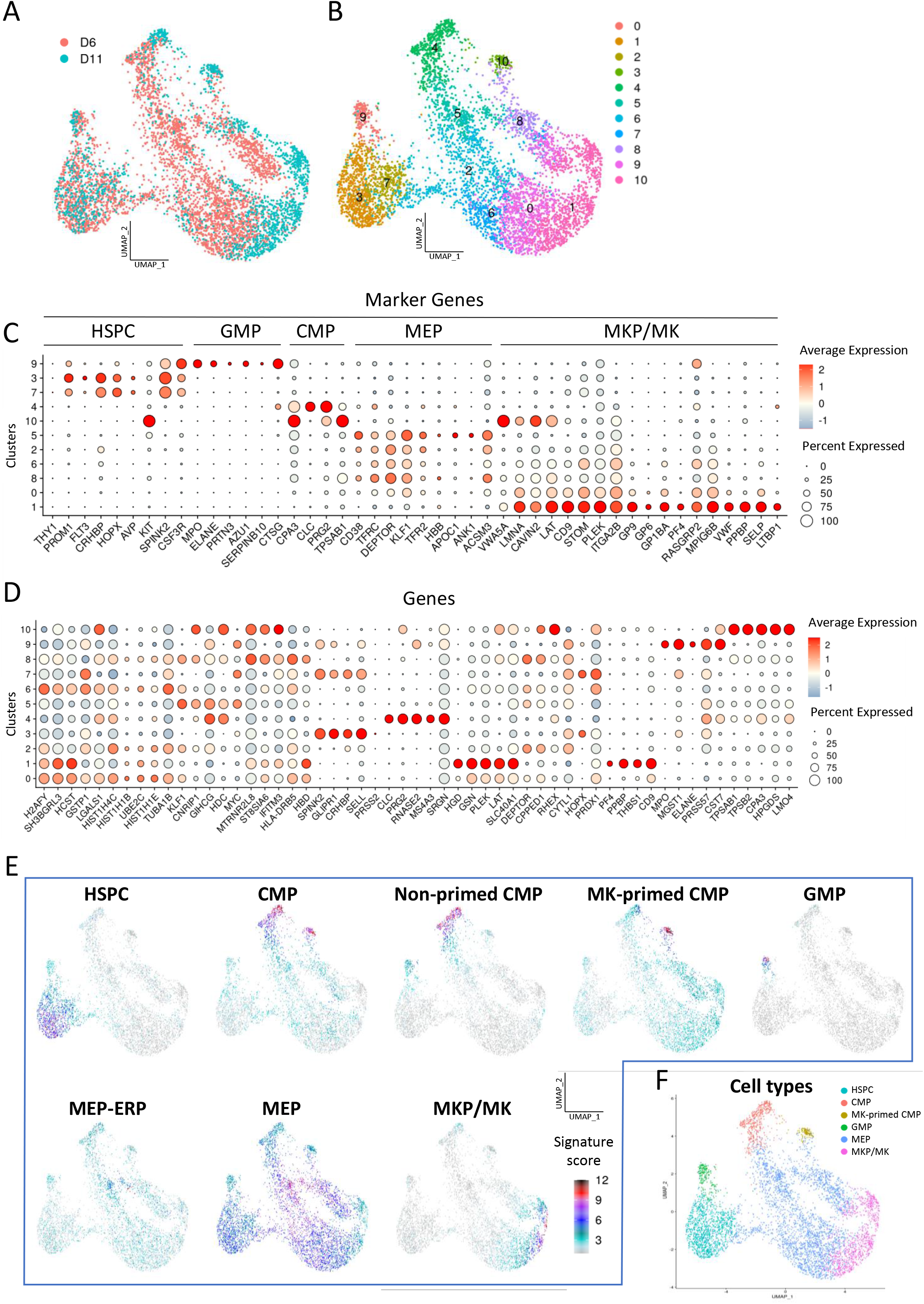
Single-cell RNA sequencing of CD34^+^ cell-derived MK-induced cells from patients with a *ETV6* variant. A- UMAP plot of *ETV6*-variant cells at days 6 and 11 after merging the two data sets. Cells analyzed at day 6 are displayed in orange, and cells analyzed at day 11 are displayed in blue. B- UMAP plot of cell clusters using a resolution of 0.8. Each of the 10 clusters is displayed with a distinct color. C- Seurat dot plot showing the average relative expression of several hematopoietic cell marker genes in each cluster. D- Seurat dot plot of the top five differentially expressed genes by cluster. Dot size represents the percentage of cells expressing the gene of interest, while dot color represents the scaled average gene expression (negative average expression indicates expression levels below the mean). E- Feature plot of the expression profile of lineage signature gene sets. F- UMAP plot of hematopoietic cells derived from *ETV6*-variant CD34^+^ cells color-coded by cell type.

Using unsupervised clustering, a total of 11 clusters were identified (Figure 3B). The cell types found in controls were also observed in patients (Figures 3C-D, Supplemental figures 6 to 8, Supplemental table 5). Clusters 3 and 7 corresponded to HSPC; cluster 4 corresponded to CMP; cluster 10 corresponded to MK-primed CMP; cluster 9 corresponded to GMP; clusters 5, 2, 6, 8 and 0 corresponded to MEP; and cluster 1 corresponded to MKP/MK. No platelet clusters were observed. The proposed lineage signatures (Supplemental table 2) were computed and displayed via feature plot, and cell types were assigned (Figures 3E-F).

### Developmental trajectory of megakaryocyte differentiation

We then assessed the single-cell transcriptome for pseudotemporal ordering of differentiation states during megakaryopoiesis in controls and patients. Using Slingshot, we inferred differentiation trajectories in cells from controls and *ETV6*-variant patients. For each condition, we assessed the overall trajectory structure of each lineage (rooted tree) by generating the transcriptomic distance matrix using the manually set root-cluster (HSPC; cluster 10 for controls and cluster 3 for patients).

Several lineages were identified in each condition (5 and 4, for control and patients, respectively) (Supplemental figure 9). We observed a lineage of MK differentiation (Figures 4A-B), which bypassed the CMP cell type and differentiated directly from HSPC into MEP. In this MK trajectory, stem-cell markers such as *CD34, CD38* and *HLA-DRA* decreased, while the MK markers *GP9* and *PF4* displayed increased expression (Figure 4C).

During differentiation from HSPC to MK, the TF *GATA1, FLI1* and *TAL1* displayed increased expression in control cells, while *GATA2* displayed high expression levels in immature cells and reduced expression in later stages. During differentiation, *ETV6* and *RUNX1* expression levels were low and stable (Figure 4D, Supplementary figures 3 and 7).

**Figure 4.**
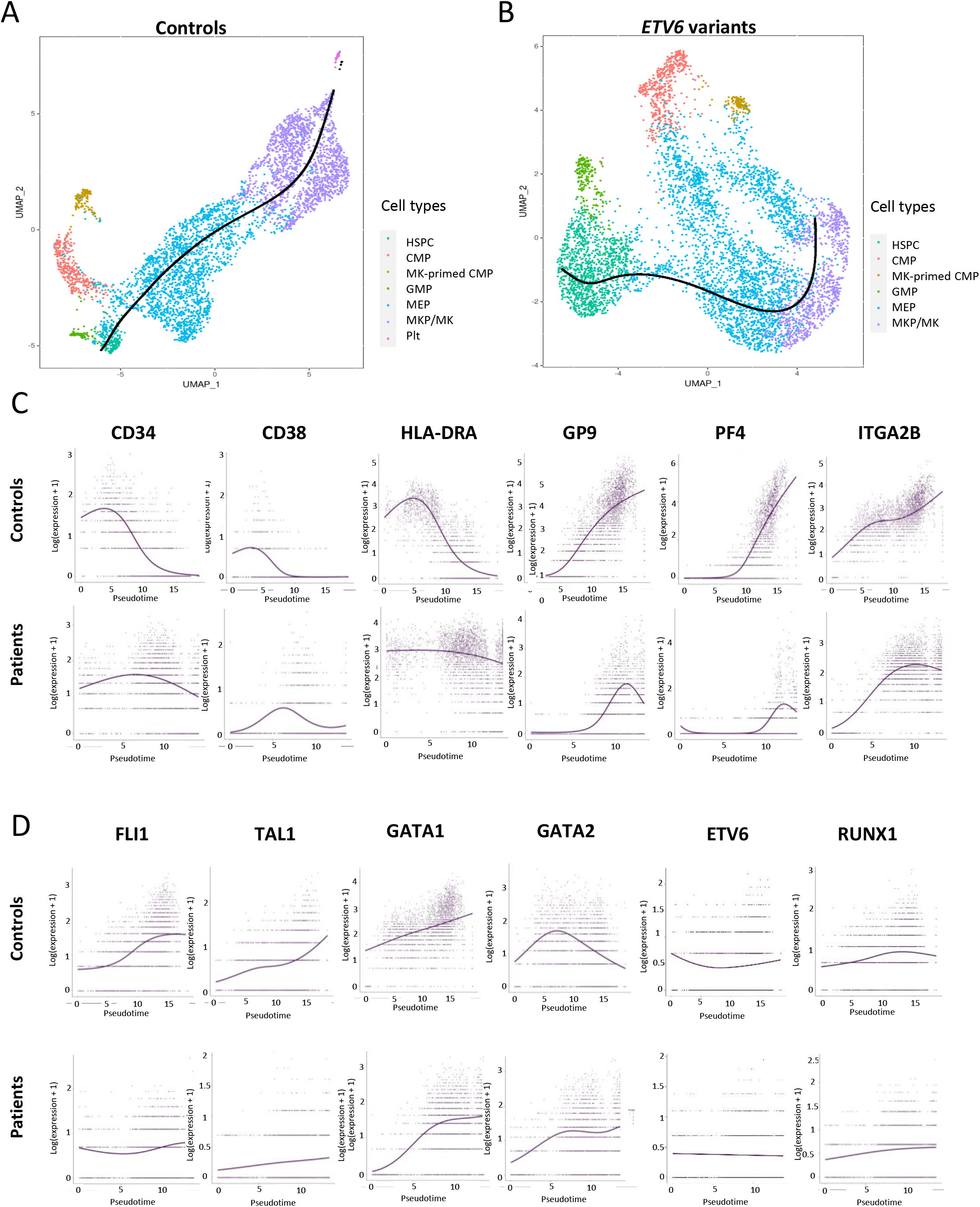
Megakaryocyte differentiation in healthy controls and *ETV6* patients. A- Pseudotime ordering was performed using Slingshot to reconstruct the hierarchical structure of hematopoietic cells derived from healthy controls. Cells are color-coded by type (HSPC, CMP, MK-primed CMP, GMP, MEP, MKP/MK and platelets (Plt)). B- Pseudotime ordering was performed using Slingshot to reconstruct the hierarchical structure of hematopoietic cells derived from *ETV6* patients. Cells are color-coded by type (HSPC, CMP, MK-primed CMP, GMP, MEP and MKP/MK). C- Dynamic expression of immature and differentiation markers along the differentiation trajectory from HSPC to MK in healthy controls (upper panel) and *ETV6* patients (lower panel). D- Dynamic expression of transcription factor genes along the differentiation trajectory from HSPC to MK. The x-axis shows pseudotime estimated by fitGAM, while the y-axis shows normalized gene expression.

Compared with controls, *ETV6*-deficient cells displayed stable expression of *CD34, HLA* (*HLA-DRA* (Figure 4C) and *HLA-DPB1, HLA-DPA1, HLA-DRB5, HLA*-*DRB1, HLA*-*A, HLA*-*E, HLA*-*DMA, HLA*-*C*, HLA-B, *HLA*-*DQA2, HLA*-*DQB1* and *HLA*-*DQA1* (data not shown)) and *GATA2* during MK differentiation; these genes remained highly expressed at the MKP/MK stage (fold change (FC) ETV6 vs. WT) = 2.5 (*CD34*), 1.5 to 8.8 (*HLA*), 1.6 (*GATA2*), adjusted p-value <0.05). The expression levels of *GATA1, FLI1* and *TAL1* were lower than that observed in controls. We observed a delayed response time for *CD38, GP9* and *PF4* (Figures 4C-D). These results suggest that *ETV6*-variant carriers exhibit a delay in differentiation.

### Aberrant populations in *ETV6*-variant cells identified via single-cell RNA sequencing

The full data set (control and patients together) was analyzed to compare controls against patients. The cell types identified in control and patient data sets were transferred to the full data set using the cell identity barcode (Figure 5A). The day 6-11 results and the differences between control and patient for each cell type are shown in Figure 5B-C. For the early stages (HSPC, CMP and GMP), the transcriptome profiles of controls and patients were markedly similar. By contrast, distinct gene expression patterns were observed for the MEP and MKP/MK populations. Cell type distribution differed between controls and patients (Figures 5D and E). Patients harboring *ETV6* variants displayed an increased proportion of HSPC (17±6.9 vs 3.0±0.7%) and a decreased proportion of MKP/MK (16.9±4.1 vs 38.5±1%) compared with controls. Furthermore, the MKP/MK populations in patients were characterized by a reduced number of detected genes (mean±SD: 2,945±927 vs 4,884±1,112, p<0.0001) and RNA count (mean±SD: 12,444±6,051 vs. 29,960±14,522, p<0.0001). Overall, these results indicate that *ETV6* mutations perturb megakaryopoiesis, resulting in a delay in cell differentiation and the development of aberrant MEP and MK populations.

**Figure 5.**
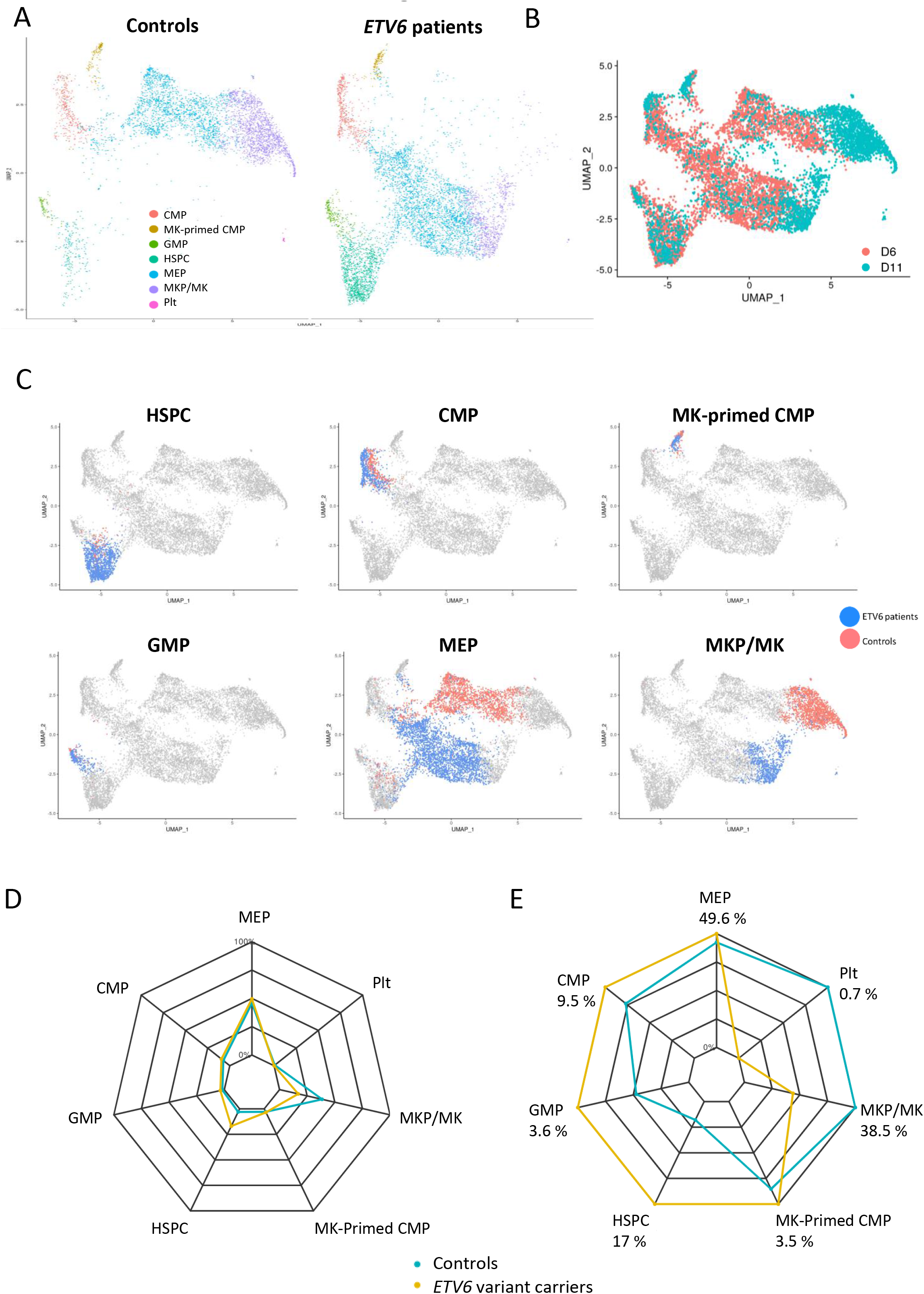
Single-cell RNA sequencing of cells derived from *ETV6*-variant carriers and healthy controls. A- UMAP plot of hematopoietic cells derived from control and *ETV6*-variant CD34^+^ cells at days 6 and 11. Cells are color-coded by cell type (HSPC, CMP, MK-primed CMP, GMP, MEP, MKP/MK and platelets (Plt)). B- UMAP plot of hematopoietic cells derived from control and *ETV6*-variant CD34^+^ cells at days 6 and 11. Cells analyzed at day 6 are indicated in orange, and cells analyzed at day 11 are indicated in blue. C- UMAP plot showing one cell type per plot. *ETV6* patient cells are indicated in blue, and control cells are indicated in orange. D- Radar plot showing the percentage of cell types for each genotype (*ETV6* patients = yellow line; control volunteers = blue line). E- The same radar plot shown in Figure 5D with a zoom on maximal values. The maximal value is indicated for each cell type.

### Patient samples displayed highly modified regulon activity and functional aberrant cell populations

We applied the SCENIC workflow to full dataset to better characterize the aberrant MEP and MKP/MK populations observed in the *ETV6* patients. The UMAP reduction, with the 312 combined regulon activity scores in each cell according to transcriptome signature designation (cell type) is shown in Supplemental figure 10A. We observed differences restricted to the MEP and MKP/MK stages. To further analyze this defect, the activity of individual regulon was also evaluated in all cells (Supplemental figure 11). Compared to the initial stages (HSPC, GMP, CMP and MK-primed CMP), we observed marked differences at the MEP and MKP/MK stages between patients and controls. The top 20 differentially active regulons are shown in Supplemental Figure 12. Common regulons were observed at the two cell stages (e.g., MAX, SPI1, GATA2, IRF5). Differences in regulon activities were more pronounced at the MKP/MK stage. The activity of 12 remarkable regulons in each cell type is presented in Supplemental figure 10B. For the latter, hyperactivity was observed in patient cells. Overall, these results complement the previous observations and indicate that *ETV6*-variants play a functional role in MEP and MKP/MK differentiation. Hyper or hypoactivity of every regulon is available in Supplemental Table 3.

### Deregulated pathways in *ETV6*-variant cells identified via single-cell RNA sequencing

When comparing controls and patients, the number of DEGs (adjusted p-value < 0.05) increased over the course of cell differentiation, especially at the MEP stage (HSPC, DEGs=38, 23 upregulated genes, 15 downregulated genes; CMP, DEGs=56, 29 upregulated genes, 27 downregulated genes; MK-primed CMP, DEGs=30, 15 upregulated genes, 15 downregulated genes; GMP, DEGs=32, 21 upregulated genes, 11 downregulated genes; MEP, DEGs=239, 100 upregulated genes, 139 downregulated genes; and MKP/MK, DEGs=942, 339 upregulated genes, 603 downregulated genes) (Figure 6A). In MEP and MKP/MK populations, DEGs were enriched in various biological pathways in multiple gene set databases (GO Biological process (Figures 6B-C), KEGG (Supplemental figure 13) and Reactome (Supplemental figure 14). Top 10 GO deregulated pathways are available in Supplemental table 6 and 7. We observed downregulation of “mitochondrial”, “mRNA processing, splicing and localization to nucleus” and “DNA repair and cellular responses to DNA damage” pathways. The “translation” pathway was among the most upregulated pathways (Supplemental table 8).

**Figure 6.**
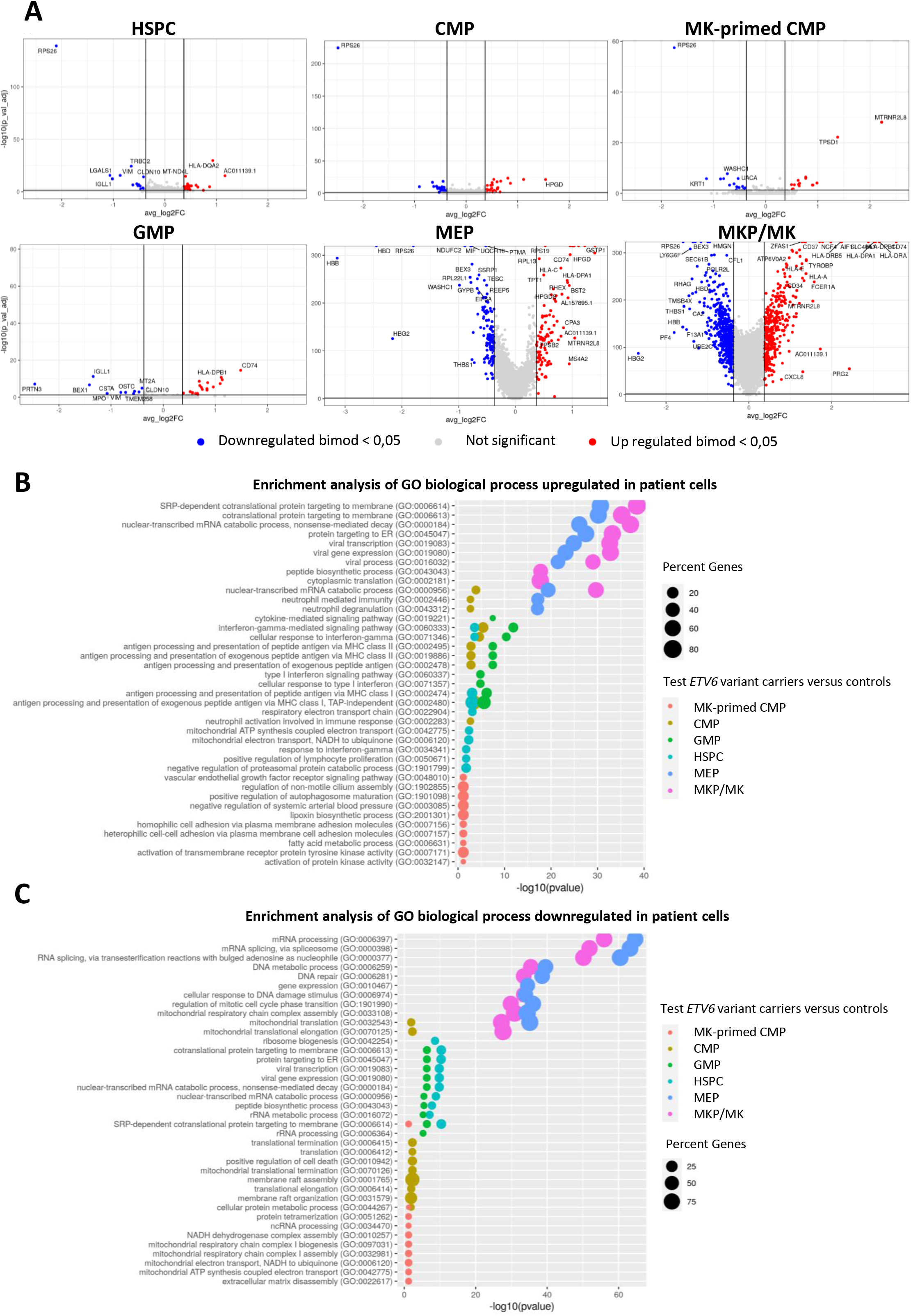
Differentially expressed genes in each cell type. A- Volcano plots of upregulated (red) and downregulated (blue) genes in *ETV6* patients vs. controls for each cell type (HSPC, CMP, MK-primed CMP, GMP, MEP and MKP/MK), FC> 1.3 [i.e., log2 FC>0.37], adjusted p-value < 0.05). B- Bubble plot of the top enriched GO biological processes based on differential gene expression by cell type (classified by p values). The upper panel shows the upregulated pathways, while the lower panel shows the downregulated pathways. Cells are color-coded by cell type (HSPC, CMP, MK-primed CMP, GMP, MEP, MKP/MK)

### Mitochondrial pathway in patients harboring an *ETV6* variant

As several mitochondria-specific processes were downregulated in *ETV6*-variant cells (i.e., mitochondrial translation, oxidative phosphorylation, respiratory electron transport chain, ATP synthesis, assembling and biogenesis complex), we further assessed the mitochondrial defects using the MitoXplorer2.0 pipeline to evaluate 36 mitochondrial functions/pathways among the DEGs^32^. The greatest FC were observed for oxidative phosphorylation, ROS defense, glycolysis, import and sorting, and mitochondrial translation (Figure 7A, Supplemental figures 15 to 17). Sixty-five of 430 downregulated mitochondrial genes contained sequences that correspond to the canonical ETV6 binding site (C/AGGAAG/A) (Normalized Enrichment Score NES 4.85, rank 389e/562, Supplemental figures 15B, Supplemental table 9).

**Figure 7.**
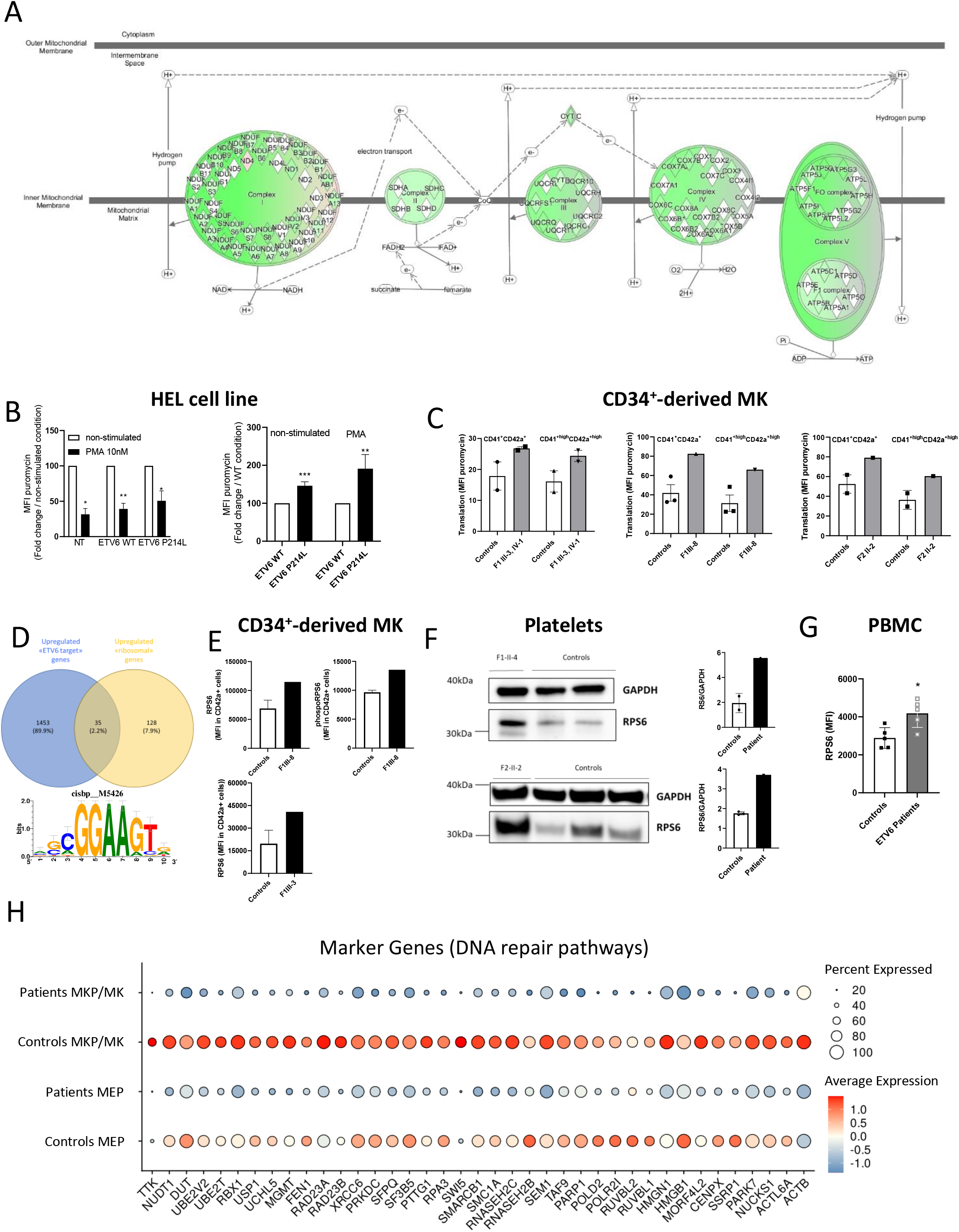
Deregulated pathways in *ETV6*-variant MKP/MK. A- Ingenuity pathway schematic diagram of *ETV6* variant-mediated gene modulation in the mitochondrial electron transport chain. The color gradient displays the gene expression fold change (green = downregulated, white = unaltered, and red = upregulated). B- Translation levels in HEL cells in basal conditions and after PMA (phorbol 12-myristate 13-acetate) stimulation (NT = non-transduced cells, ETV6 WT = cells transduced with wild type *ETV6*, and ETV6 P214L = cells transduced with the *ETV6* P214L variant). Translation levels were evaluated via flow cytometry by assessing protein synthesis after incorporation of puromycin and staining with monoclonal anti-puromycin-APC antibody. *P<0.05, **P<0.01, ***P<0.001, one-sample t-test. C- Translation levels in CD34^+^ cell-derived MK at day 11 of culture in controls and P214L variant cells (F1 III-3 and IV1, F1III-8) and day 14 for F417Lter4 variant cells (F2 II-2). Translation levels were evaluated via flow cytometry by assessing protein synthesis after incorporation of puromycin and staining with monoclonal anti-puromycin-FITC antibody. D- Venn diagram of upregulated genes with an *ETV6* binding site (blue circle) and ribosomal genes (yellow circle) in MKP/MK isolated from *ETV6-*patients vs controls. Among all upregulated genes carrying an ETV6 binding site, 35 genes were associated with translation pathways. E- Flow cytometry assessment of RPS6 and phoshoRPS6 in CD34^+^ cell-derived MK from F1III-8 and four healthy controls (analyzed at the same time, at day 14) and RPS6 CD34^+^ cell-derived MK from F1III-3 and three healthy controls (analyzed at the same time, at day 13). RPS6 was analyzed by gating on CD42a^+^ cells. F- Western blot analysis of RPS6 expression in washed platelets from healthy controls and patients harboring ETV6 variants. GAPDH was used as a protein loading control. Quantification of band intensity is shown on the right. G- Flow cytometry assessment of RPS6 in peripheral blood mononuclear cells (PBMC) from five controls and five *ETV6* patients (F1-II4, III3, III8, VI1 and F2-II2). PBMC were frozen on the day of patient consultation and then thawed and cultured for four days before RPS6 analysis (all samples were analyzed at the same time). * P<0.05, Mann-Whitney t-test. H- Seurat dot plot of the DEG in DNA repair pathways in MEP and MKP/MK cell populations (patient and control samples). Dot size represents the percentage of cells expressing the gene of interest, while dot color represents the scaled average gene expression (negative average expression indicates expression levels below the mean).

The number of DEGs progressively increased over the course of cell differentiation (Supplemental figures 15C and 16). *DNML1*, which encodes a key mediator of mitochondrial division DRP1, and *CYCS*, which encodes for cytochrome C, were downregulated in *ETV6*-variant MK (Supplemental figure 18).

### Increased translation in patients harboring an *ETV6* variant

Translation was the dominant upregulated pathway. Translation levels were evaluated in HEL cells transduced with P214L or WT *ETV6* before and after PMA (phorbol 12-myristate 13-acetate)-induced differentiation. As expected, PMA stimulation led to reduced translation in all samples (Figure 7B). Translation levels were higher in *ETV6*-deficient cells at the basal state and after PMA stimulation (Figure 7B). Compared with control cells, higher translation levels were observed in CD34^+^-derived MK isolated from three *ETV6*-carriers at day 11 (F1 III-3, III-8, IV-1) and one at day 14 (F2 II-2) (Figure 7C). *RPS* and *RPL* were the most upregulated genes, which code for ribosomal protein small subunit and ribosomal protein large subunit, respectively (Supplemental figure 19). Among these 163 upregulated ribosomal genes, 35 genes contained sequences that correspond to the canonical ETV6 binding site (NES 4.85, rank 40e/512, Supplemental table 9), including RPS6 (Figure 7D). The RPS6 kinase genes (*RPS6KA1, RPS6KA3, RPS6KC1*) also contained ETV6 binding sites (Supplemental table 9). Compared with controls, RPS6 was overexpressed in CD34^+^-derived MK (day 14) from one patient and platelets from two patients (Figures 7E-F). RPS6 was also upregulated in peripheral blood mononuclear cells (PBMC) from five patients with an *ETV6* variant compared with controls (Figure 7G).

### Downregulated DNA repair pathways in *ETV6*-variant carriers

Several pathways associated with DNA repair and cellular response to DNA damage were downregulated in patient MEP and MK (Supplemental table 8). Some genes involved in major DNA repair pathways displayed reduced expression levels: *FEN1* (base excision repair), *MGMT* (direct reversal of DNA damage), *RAD23A* and *RAD23B* (nucleotide excision repair), and *XRCC6* and *PRKDC* (non-homologous end joining)^33^. Caretaker genes indirectly involved in maintaining genomic stability (e.g., *TTK, NUDT1, DUT, UBE2V2*) were also downregulated in MK harboring *ETV6* variants (Figure 7H).

## Discussion

Using single-cell transcriptome profiling of MK cell cultures derived from peripheral CD34^+^-cells, we established a signature for each cell stages of megakaryopoiesis. Both the gene expression profiles and the regulon profiles confirmed distinct cell-type signatures for each stage of differentiation. We also observed a differentiation trajectory in which MEP developed directly from HSPC and bypassed the CMP stage. A small MK-primed CMP population was also evidenced. Cells harboring *ETV6* variants displayed significantly distinct gene expression profiles starting at the MEP stage. Our results also reveal concomitant differences in the activity of specific regulons. Furthermore, the observed dysregulation of several pathways indicates that *ETV6* deficiency affects key cellular processes associated with mitochondria function, translation and DNA repair, which may represent promising mechanistic targets.

In this study, we highlight a biased megakaryopoiesis pathway that may be due to stress-driven hematopoiesis as a result of the culture conditions^34^. HSPC directly gave rise to MEP without passing through the CMP stage, as previously suggested^35^. Furthermore, it has been hypothesized that MK-biased hematopoietic stem cells (HSCs) represent the hierarchical apex with MEP developing directly from HSCs^36^, although this view has been challenged. We did not observe a MK-biased HSPC population^37,38^ either because (1) the analysis time points (days 6 and 11) were too late to detect these cell types, (2) clustering was insufficient to visualize this small population, or (3) mRNA did not enable detection of MK-biased HSPC, as previously reported^39^. However, we identified a MK-primed CMP population that co-expressed megakaryocytic genes and displayed high *KIT* expression levels (27% of the CMP population). High KIT expression levels represented a good marker of MK-biased CMP. Previous murine *in vitro* and *in vivo* functional studies have demonstrated that HSCs with higher levels of c-Kit signaling preferentially differentiate into MK^40^. In human bone marrow, MK-primed CMP likely represented the major megakaryopoiesis pathway independent of the canonical MEP lineage^41^. This *in vitro* model may be a valuable tool to investigate these biased pathways.

We characterized each cell type by analyzing the activity of TF specific to each cell stage, which confirmed the gene expression designations. HSPC were characterized by the expression of the regulons TCFL2, TCF and HOXA9, which are involved in stem cell maintenance^42–44^. C/EBP family members are critical for myelopoiesis^45,46^. Accordingly, we observed noticeable C/EBP activity at the CMP and GMP stages. Interestingly, we observed activated AhR in the MK-primed CMP population, thus indicating that AhR modulation plays a role in MK maturation, as previously described^47–49^. We also observed KLF1 activity in MEP, as KLF1 is involved in MEP lineage decisions and commitment^50^. Finally, NFE2 and GATA1 were expected markers of MK^51,52^. Overall, these findings demonstrate that characterizing TF activity at the single-cell level can be useful to phenotype cell types.

Applying the lineage signature used in control cells, we found that *ETV6*-deficient cells displayed the same cell type with a higher proportion of HSPC, a lower proportion of MK and an absence of platelets, which correlates with the thrombocytopenia phenotype observed in patients harboring an *ETV6* variant. We detected marked differences in the TF regulatory network; the most-affected regulons displayed higher activity levels in MEP and MKP isolated from patients. This finding is in accordance with a loss of ETV6 repressor activity, which plays a role in the development of the pathology observed in patients. Also, some regulons hypoactivity was detected, challenging the unique repressor role of ETV6. It may be both an activator and a repressor depending of the genomic context and its co-factors, making it a potential pioneer TF like SPI1^53^.

Analysis of several gene enrichment databases displayed significant differences. Assessing the ten highest scores and after grouping the pathways with common genes, we observed deregulation of pathways linked with mitochondria function, translation and DNA repair in MEP and MKP/MK.

Our results indicate that patients harboring an *ETV6*-variant display decreased expression of genes involved in mitochondrial metabolism. We observed a marked decrease in the expression levels of *DNM1L*, which encodes DRP1 and has been shown to enhance the production of proplatelet forming MK^54,55^. These findings suggest that the mitochondria plays a role in ETV6-RT development, which merits further functional analysis.

Translation was the most upregulated pathway observed in MEP and MK patient cells. Altered expression levels of numerous ribosomal proteins (RP) genes were observed, such as downregulation of *RPS26* and upregulation of more than 30 RP with a FC>1.4. Very recent data has emphasized the unexpected contribution of ribosomal biogenesis in hematopoiesis and megakaryopoiesis. Treatment of mice or humans with the ribosomal biogenesis inhibitor CX-5461 results in an increase in circulating platelets, which is associated with the platelet/MK-biased hematopoietic pathway^56^. Additionally, ribosome biogenesis is involved in mediating the transition between proliferation and differentiation of erythroid progenitors^57^. GTPase-Dynamin-2 deletion in platelets and MKs induce a severe thrombocytopenia and bleeding diathesis in mice and result in upregulation of genes involved in ribosome biogenesis in erythroblast^58^. Several ribosomal proteins have been found to play a role in extra-ribosomal functions, including induction of apoptosis, tumor suppression, regulation of development, and DNA repair^59^. Our results highlight a significant decrease in DNA repair pathways in patients. Overall, these findings indicate that *ETV6* mutations are responsible for defects in translation and DNA repair pathways, which may contribute to leukemia predisposition.

Increased translation was confirmed at the functional level. Protein synthesis was increased in CD34^+^ cell-derived MK in patients and hematopoietic cell lines transduced with an *ETV6* variant. As *RPS6* contains putative ETV6 binding sites, which was confirmed via ChIP-sequencing (UCSC Genome Browser on GSM2574795, GSM2534228, GSM2574796), and *Rps6*-deficient mice display features of megakaryocytic dysplasia with thrombocytosis, this gene likely plays a role in thrombopoiesis ^60,61^. Remarkably, total RPS6 antigen levels were increased two-fold in patient MK, platelets and PBMC compared with controls. Indeed, further investigation is required to elucidate the specific role that RPS6 plays in ETV6-RT.

In summary, our study demonstrates the heterogeneity of human CD34^+^ cell-induced MK differentiation *in vitro*, a novel MK-primed CMP population, and a major differentiation trajectory in which HSPC develop directly into MEP and bypass the CMP stage. We found that *ETV6* variations cause defects in early hematopoietic stages and result in an aberrant MEP and MK populations with deregulated translation and DNA repair pathways. These findings provide novel insight into megakaryopoiesis and *ETV6* function that may be applied to develop targeted therapeutic strategies to alleviate platelet defects.

## Acknowledgments

This work was supported by Aix-Marseille University (AMIDEX “Emergence et innovation” ngSUMMIT), the Agence Nationale de la Recherche (JCJC MOST) and the Institut National de la Santé et de la Recherche Médicale (PIA Biofit). The authors acknowledge the members of the French Reference Center for Inherited Hereditary Platelet Disorders for their contribution regarding clinical analyses and the GBiM platform for the sequencing and discussion.

## Authorship contributions

TB, LH and DP performed bioinformatic analyses. EG, EA, VS and DB performed the culture and functional experiments. CL, MIK, ML and PS performed the clinical and biological characterization of patients. DPB and MCA conceived and supervised the project. DP and MP directed the project, designed the study, analyzed the data and wrote the manuscript.

## Disclosure of conflicts of interest

The authors have declared that no conflict of interest exists.

## Notes

https://zenodo.org/record/6980009

### Competing Interest Statement

The authors have declared no competing interest.

