## Supplemental Material 1 for "Single-cell analysis of megakaryopoiesis in peripheral CD34^+^ cells: insights into ETV6-related thrombocytopenia"

#### Methods

##### Patient enrollment and DNA sequencing

Patients and healthy volunteers were recruited at the Center for the Investigation of Hemorrhagic and Thrombotic Pathologies at Marseille University Hospital. Volunteers were included in the study after informed written consent was obtained, in accordance with protocols approved by the national review process (authorization number 20200T2-02). Two high-throughput sequencing methods were used to identify *ETV6* variants as previously described[1,2]. Two *ETV6*-variants were evaluated in this study, the novel variant pF417LTer4 (NM_001987, c.1251del) and a previously described variant p.P214L (NM_001987, c.C641T).

##### *In vitro* megakaryocyte differentiation

After density gradient separation (Eurobio), circulating $CD34^+$ cells were purified using positive selection with magnetic beads (Miltenyi-Biotec) and then cultured in StemSpan Serum-Free Expansion Medium II supplemented with Megakaryocyte Expansion Supplement (Stemcell Technologies) as previously described[2] (Figure 1A).

##### Cell preparation and single-cell RNA sequencing

$CD34^+$ differentiated cells from two healthy controls and two patients harboring *ETV6* variants were harvested from culture at days 6 and 11 (Figure 1A). Cell concentrations and viability were assessed after trypan blue staining with an automated cell counter (EveTM NanoEntek). The cell samples from each volunteer were labeled with a distinct hashtag oligo (TotalSeq, Biolegend) and pooled. Single-cell isolation was then carried out with the 10x Genomics Technology using the Chromium Next GEM Single Cell 5' Kit v2 (ref 1000263) according to the manufacturer's protocol. Single-cell cDNA synthesis and sequencing libraries (days 6 and 11) were prepared with single-cell 5' Library and Gel Bead kit. Libraries were sequenced applying a 75-bp paired end reads format with a Next-seq500 (GBiM platform) (parameters, read 1: 26 cycles, i7: 8 cycles, read 2: 57 cycles).

##### Single-cell RNA sequencing analysis

mRNA library reads were aligned to the hg38 reference genome (GRCh38-2020-A) and quantified using the cellranger count pipeline (v5.0.0). For cell hashing, antibodies were quantified using CITE-seq-count (v1.4.3) with default parameters. The resulting mRNA and hashtag oligo data matrices were imported into R (v4.1.2), and downstream analyses were performed using the Seurat package (v4.0.5).

**Analysis of single-cell RNA sequencing data**

For data preprocessing, mRNA library reads were aligned to the hg38 reference genome (GRCh38-2020-A) and quantified using the cellranger count pipeline (version 5.0.0). For cell hashing, antibodies were quantified using CITE-seq-Count (version 1.4.3) with the default parameters. The resulting mRNA and hashtag oligo data matrices were imported into R (version 4.1.2), and downstream analyses were performed using the Seurat package (version 4.0.5). Cells were demultiplexed using the HTODemux and MULTISeqDemux functions. To enhance demultiplexing, we identified specific single nucleotide variants for each sample using the Souporcell method (version 2.0). Applying both techniques, cell multiplets and negatives were removed from the filtered cellranger barcode matrix. Before mRNA analysis, we filtered out low quality cells (i.e., cells with an mRNA count <1000 and cells with less than the 8% threshold of mitochondrial gene expression, which was calculated using the scater::isOutlier function) and genes expressed in few cells (e.g., genes expressed in less than 3 cells). Expression raw counts were then normalized and scaled using the Seurat NormalizeData and ScaleData functions, respectively. Samples from the two time points (day 6 and day 11) were merged using the Seurat merge function. Integration by fixing anchors was not necessary, the merge quality and absence of batch effects were subsequently assessed by overlapping several cell groups at both time points. After normalization and regression scaling (based on cell cycle genes), two-dimensional reduction was performed. The merged object was first reduced using principal component analysis (PCA) (Seurat::RunPCA) with the 1000 most variable genes (Seurat::FindVariableFeatures). The PCA reduction was used to generate a UMAP plot with the first 20 components (Seurat::RunUMAP). The nearest-neighbor graph (*k*-nn) was generated using the Seurat::FindNeighbors function with a k parameter equal to 30 and the first 10 components. Cell clusters were identified using the Seurat FindClusters function with the Louvain algorithm (with a resolution of 1.2 for healthy controls and 0.8 for *ETV6*-variant carriers). Based on the expression of *CD19* and *MS4A1* (B cells), *CD3* (T cells) and *ITGAM* (monocytes/macrophages), we excluded off-target clusters from our analysis. The Seurat analysis was performed on this new data subset using 1,000 variable genes, default parameter scaling, nearest-neighbors graph with 10 components and k parameter equal to 30, clustering resolution of 0.8 and UMAP with 28 components. This Seurat analysis was then split between controls and *ETV6* patients. Each subset object was individually processed as previously mentioned. Based on published hematopoietic gene markers and the top 20 cluster-specific differentially expressed genes, we established cell type-specific signatures, which were scored on each cluster for controls and patients independently and then assigned each cluster to a cell type. We transferred these designations onto the merged object. The differential gene expression analysis was performed for each cell type (controls vs. *ETV6* patients) with the log fold-change threshold equal to 0 and using the non-parametric Wilcoxon test. After filtering differentially expressed genes with a p-value < 0.05 and either average log2 fold change > 0 or average log2 fold change < 0, enrichment analysis was performed on five different databases (Transfac and Jaspar, KEGG 2019, GO Biological process 2018, Reactome 2016 and Transcription factor PPI) and summarized in bubble plots. To infer overall single-cell trajectory (and therefore distinct differentiation lineages) and pseudotime variables in each condition, we used Slingshot (2.2.0) with the default parameters. To infer single-cell trajectory-based differential expression and therefore assess gene expression over pseudotime, we used tradeSeq (1.7.07) with the default parameters. To infer gene regulatory networks implicated in *ETV6*-related transcriptome dysregulation and failed megakaryocyte maturation that leads to thrombocytopenic disorders, we used the SCENIC method using a cellranger-filtered raw unique molecular identifier (UMI) matrix. All statistical tests comparing patients against controls were either Wilcoxon or bimodal tests with p value < 0.05.

**Data and code availability**

FASTQ raw data are available in the Sequence Read Archive (in progress). Processed data can be accessed from the NCBI Gene Expression Omnibus database (GSE206089). The human transcriptome reference used in our study is available on 10XGenomics website (https://cf.10xgenomics.com/supp/cell-exp/refdata-gex-GRCh38-2020-A.tar.gz) or Zenodo (https://zenodo.org/record/6980009/files/?download=1). Docker images and detailed scripts used for preprocessing and further analysis are available at Zenodo (https://zenodo.org/record/6980009) and github (https://github.com/poggiteam/ETV6_2020), respectively.

**i-cisTarget analysis**

i-cisTarget (https://gbiomed.kuleuven.be/apps/lcb/i-cisTarget/index.php) was used to predict regulatory features and cis-regulatory modules. The analysis was performed on HUGO gene nomenclature committee genes that were overexpressed or underexpressed in patient cell types, using the entire motif collection (i.e., all position weight matrix databases available). Enrichment was evaluated using human genome hg19 as a reference and gene annotation RefSeq r45 (version 6.0). Region mapping was set between 10 kb upstream and 10 kb downstream of transcription start sites. The normalized enrichment score threshold

was set at 3.0, and all other optional parameters were set by default. For each enriched motif of interest that was over the normalized enrichment score of 3.0, we analyzed the candidate target genes.
Venn diagram were performed for deregulated mitochondrial and ribosomal genes in *ETV6*-variants carrier MKP/MK. It shows the overlap of deregulated transcripts containing sequences that correspond to the canonical ETV6 binding site and deregulated genes of mitochondrial and ribosomal gene pathways.

**Site-directed mutagenesis and luciferase assays**

*ETV6* cDNA was ligated into a pcDNA3 expression vector, and mutagenesis was performed using the GENEART® Site-directed Mutagenesis System kit (Life technologies) as previously described.
Transcriptional regulatory properties of wild type (WT) and mutant ETV6 was analyzed using the luciferase reporter systems p(E74)3tk80Luc plasmid containing the luciferase gene driven by an enhancer/promoter cassette composed of three tandem copies of the ETS binding site (Lopez et al. 1999). GripTite™ 293 macrophage scavenger receptor (MSR) cells (a genetically engineered human embryonic kidney 293 cell line, Thermofisher Scientific) were transfected using Polyjet reagent (SignaGen) as specified by the manufacturer. pGL4-hRLuc plasmid was used to normalize for transfection efficiency. Cell lysates were prepared 48 hours after transfection and assayed for luciferase activity using the Dual-Luciferase® Reporter Assay system (Promega).

**Western blot**

To isolate nuclear and cytoplasmic subcellular fractions from protein lysates, GripTite 293 MSR cells were lysed 48 hours after transfection (PolyJet In Vitro DNA Transfection Reagent, SignaGen Laboratories). Fractionation was performed using NE-PER Nuclear and Cytoplasmic Extraction Reagent (Pierce). Total washed platelet (as previously described[3]) proteins and proteins from the subcellular fractions were separated on NuPAGE gels with MES SDS running buffer (Thermofisher Scientific) and transferred onto a polyvinylidene fluoride membrane. The membranes were blocked and labeled overnight with the following primary antibodies: goat anti-ETV6 (Santa Cruz Biotechnology, sc-8546), rabbit anti-lamin-B1 (Abcam, ab16048), rabbit anti-RPS6 (Cell Signaling, 2217), and mouse anti-GAPDH (Millipore, MAB374). The membranes were then incubated with the appropriate horseradish peroxidase–conjugated secondary antibody (Biorad). Chemiluminescence was then assessed using the CCD camera ImageQuant LAS 4000 (GE Healthcare) and quantified using ImageJ software (National Institutes of Health, Bethesda, MD, USA).

**Epifluorescence microscopy**

Forty-eight hours after transfection, GripTite 293 MSR cells were fixed in 4% paraformaldehyde for 20 minutes at room temperature. The cells were then washed, permeabilized with 0.3% Triton X100 in phosphate-buffered saline (PBS) for 5 minutes, blocked with 3% bovine serum albumin (BSA) PBS, and incubated overnight with rabbit anti-ETV6 antibody (Santa Cruz Biotechnology, sc-8546). Next, the cells were incubated with anti-rabbit Alexa 546-labeled secondary antibody (Life technologies, A11010), and rhodamine phalloidin (cytoskeleton) staining was performed. Finally, after washing the cells, the slides were mounted with DAPI-Fluoromount and examined using an AXIO Imager M1 microscope (Carl Zeiss, Germany).

**Lentiviral particle production and transduction**

WT and p.P214L *ETV6* DNA were subcloned into a third generation of HIV-derived lentiviral vector pRRLsin-PGK-IRES2-eGFP-WPRE (Genethon), and lentiviral stocks were prepared as previously described[4–6]. HEL (Human Erythro-leukemia) cells ($1.10^5$) were infected with lentiviral particles. After 4 hours, lentiviral particles were diluted. After 8 hours, cells were washed and cultured with RPMI-1640 medium (Gibco, Invitrogen Corporation, Carlsbad, CA, USA) supplemented with 10% FBS (Thermo Fisher Scientific, Waltham, MA, USA) and 1% penicillin/streptomycin (Gibco, Invitrogen Corporation, Carlsbad, CA, USA).

**Analysis of translation levels**

Puromycin, an aminonucleoside Tyr-tRNA mimetic antibiotic that enters the A site of ribosomes and is incorporated into nascent chain, was used to evaluate translation levels in HEL and $CD34^+$-derived megakaryocytes using ad hoc antibodies and flow cytometry[7]. After puromycin treatment (10 mg/ml, 45 min, 37°C), cells were washed in cold PBS and stained with a combination of Fc receptor blocking reagent and fluorescent cell viability markers. Cells were then treated with primary conjugated antibodies against surface markers for 25 minutes at 4°C in PBS 1X 5%FCS, 2 mM EDTA (FACS wash buffer). After washing, cells were fixed and permeabilized using FOXP3 fixation and permeabilization buffer (Thermofisher eBioscience) following the manufacturer’s instructions. Intracellular puromycin staining was performed by incubating cells with anti-puromycin monoclonal antibody labeled with Alexa Fluor 647 diluted in permeabilization buffer for 1 hour at 4°C.

**Flow cytometry-based quantification of RPS6 and phosphoRPS6**

PBMC and $CD34^+$ cell-derived megakaryocytes were fixed in 1% paraformaldehyde in PBS. Next, the fixed cells were centrifuged and suspended in permeabilization buffer (0.5% Triton X-100, 2 mM Ethylene diamine tetra-acetic acid, 0.5% BSA PBS) and labeled with mouse anti-RPS6 (Cell signaling, #55594s) and rabbit anti-phosphoRPS6 (Cell signaling, #5316). After washing the cells, the data were acquired using an Accuri C6 Cytometer or Navios and analyzed with Kaluza.

**Statistical analyses**

Quantitative variables are expressed as the mean ± SEM. Experimental data analyses were performed using GraphPad Prism software (v9.3.1). Statistical differences were determined using one-way ANOVA with Kruskal-Wallis post-hoc test, Mann Whitney test or one-sample t-test when appropriate. P-value <0.05 was considered statistically significant.

**Tables**

See Excel file

**Supp. table 1: Top 20 differentially expressed genes for each cluster in control individuals.**

**Supp. table 2: Proposed gene signatures of each hematopoietic cell type.**

Genes included in HSPC, CMP, non-primed CMP, MK-primed CMP, GMP, MEP, MEP-ERP and MKP/MK signatures.

**Supp. table 3: List of regulons identified and their targets**

**Supp. table 4: Clinical characteristics of the patients harboring an *ETV6* variant.**

**Supp. table 5: Top 20 differentially expressed genes for each cluster (*ETV6*-variant carriers).**

**Supp. table 6: Top 10 downregulated pathways (Gene Ontology Biological Process) for each cell type classified on adjusted p-values.**

**Supp. table 7: Top 10 upregulated pathways (Gene Ontology Biological Process) for each cell type classified on adjusted p-values.**

**Supp. Table 8: Synthesis of deregulated pathways in ETV6 patients.** Top deregulated pathways in each cell type (adjusted p-value <0.05), *ETV6* patients vs. controls. **Supp. table 9: Results of i-cisTarget.** List A corresponds to genes containing an ETV6 binding site and downregulated in patients' MK. List B corresponds to genes containing an ETV6 binding site and upregulated in patients' MK. List C corresponds to mitochondrial genes downregulated in patients' MK. List D corresponds to ribosomal genes upregulated in patients' MK. List E corresponds to overlap between list A and C. List F corresponds to overlap between list B and D.

### Supplementary Figure 1

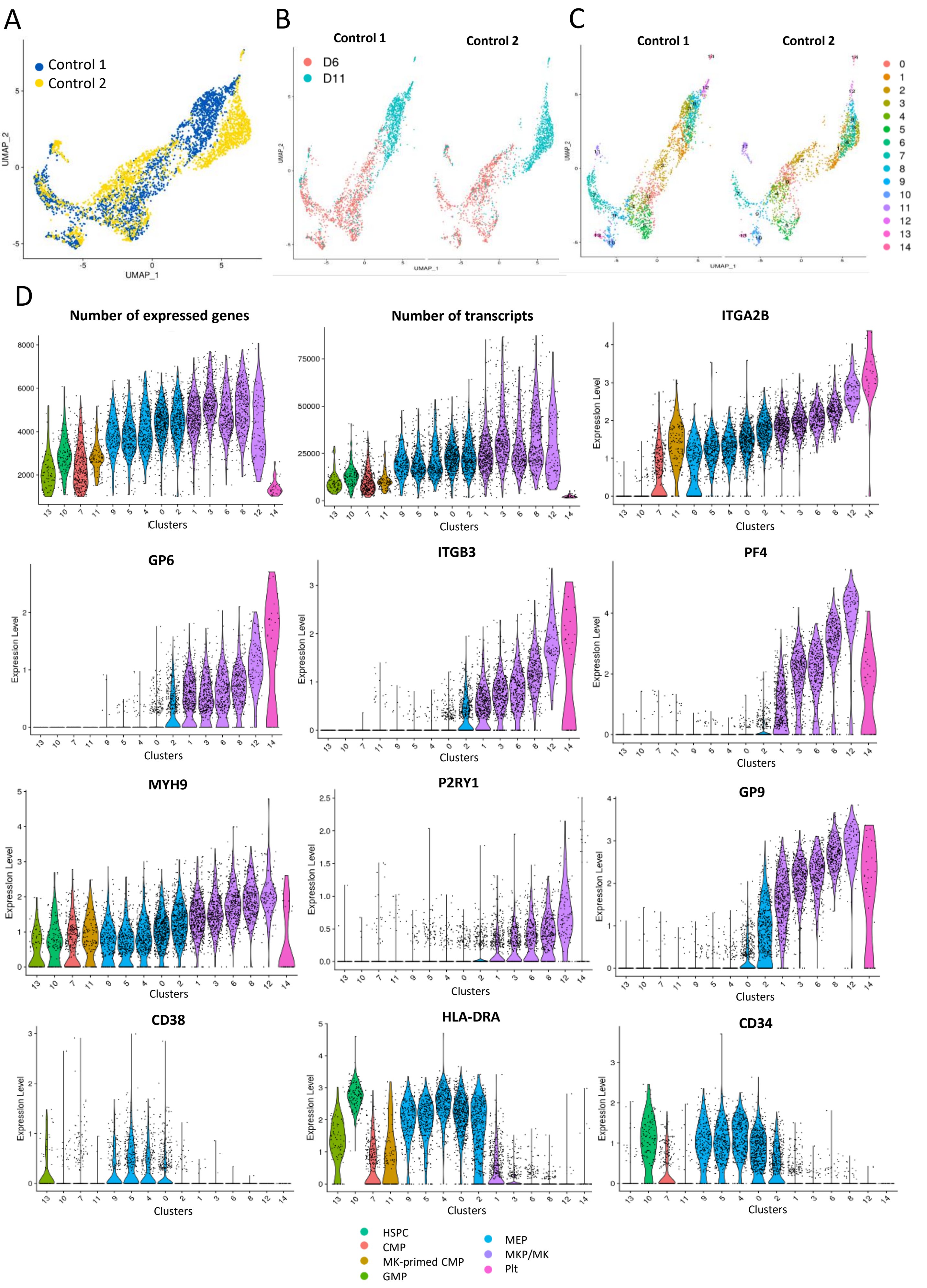

### Supplementary Figure 2

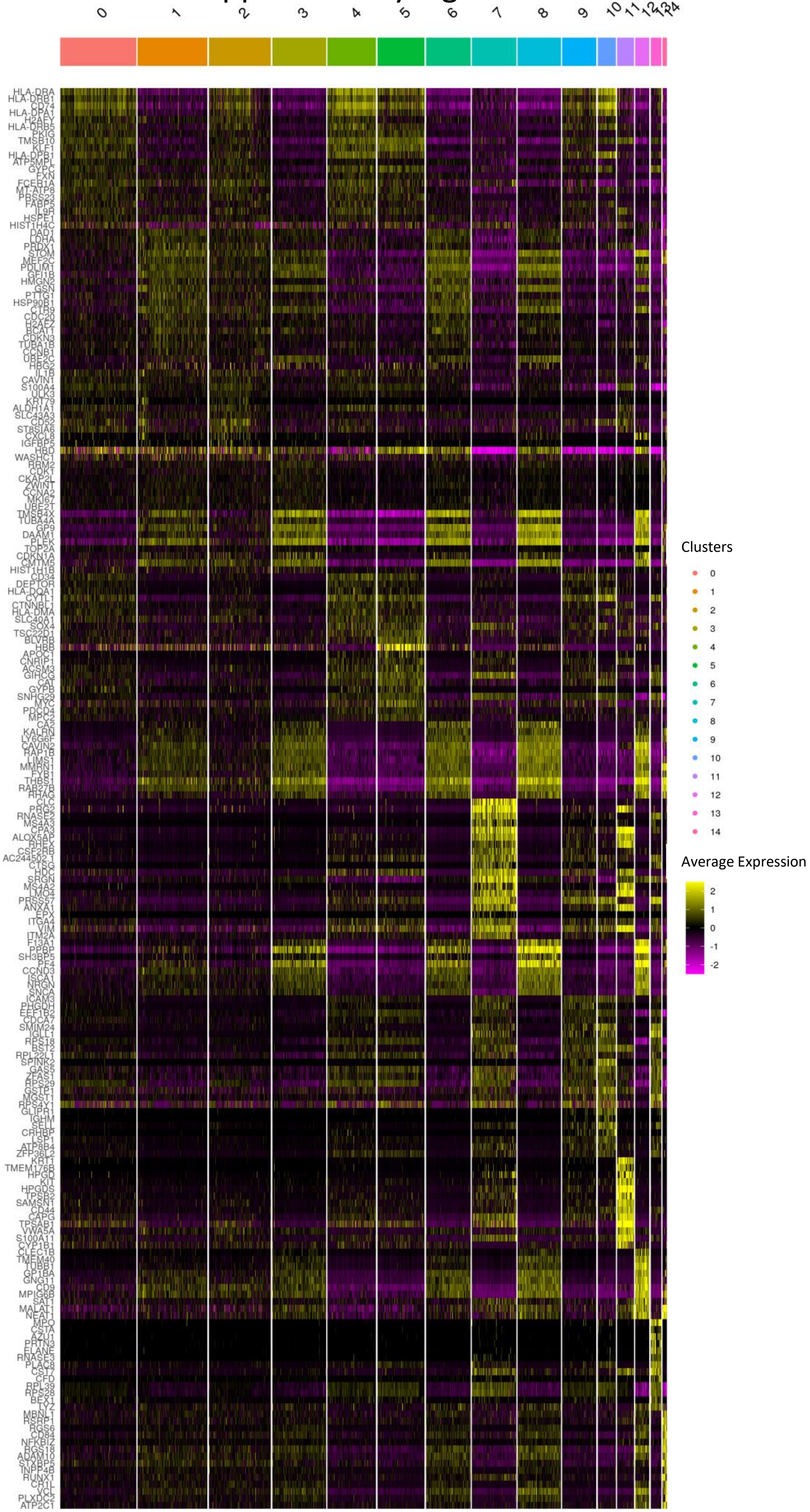

### Supplementary Figure 3

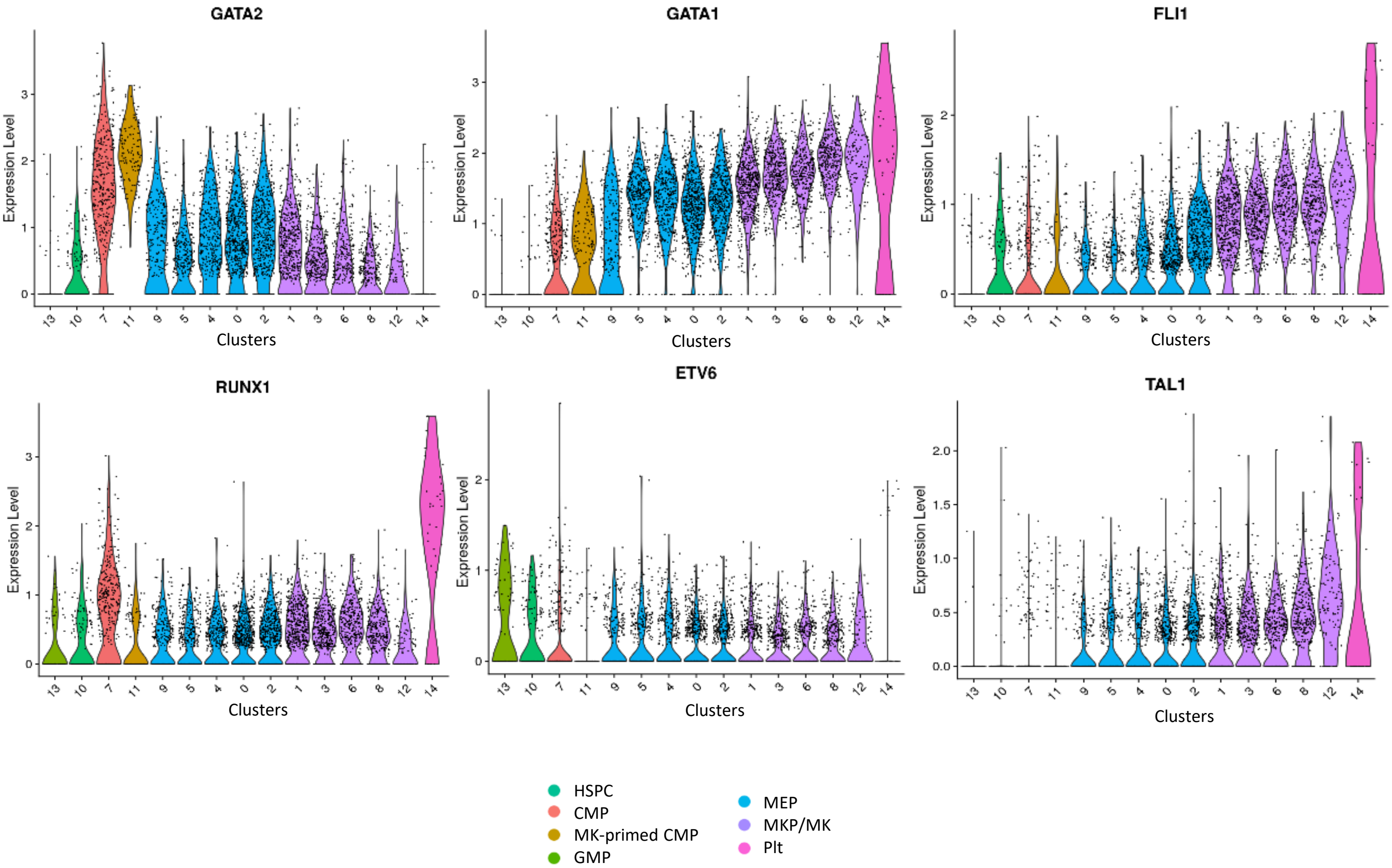

### Supplementary Figure 4

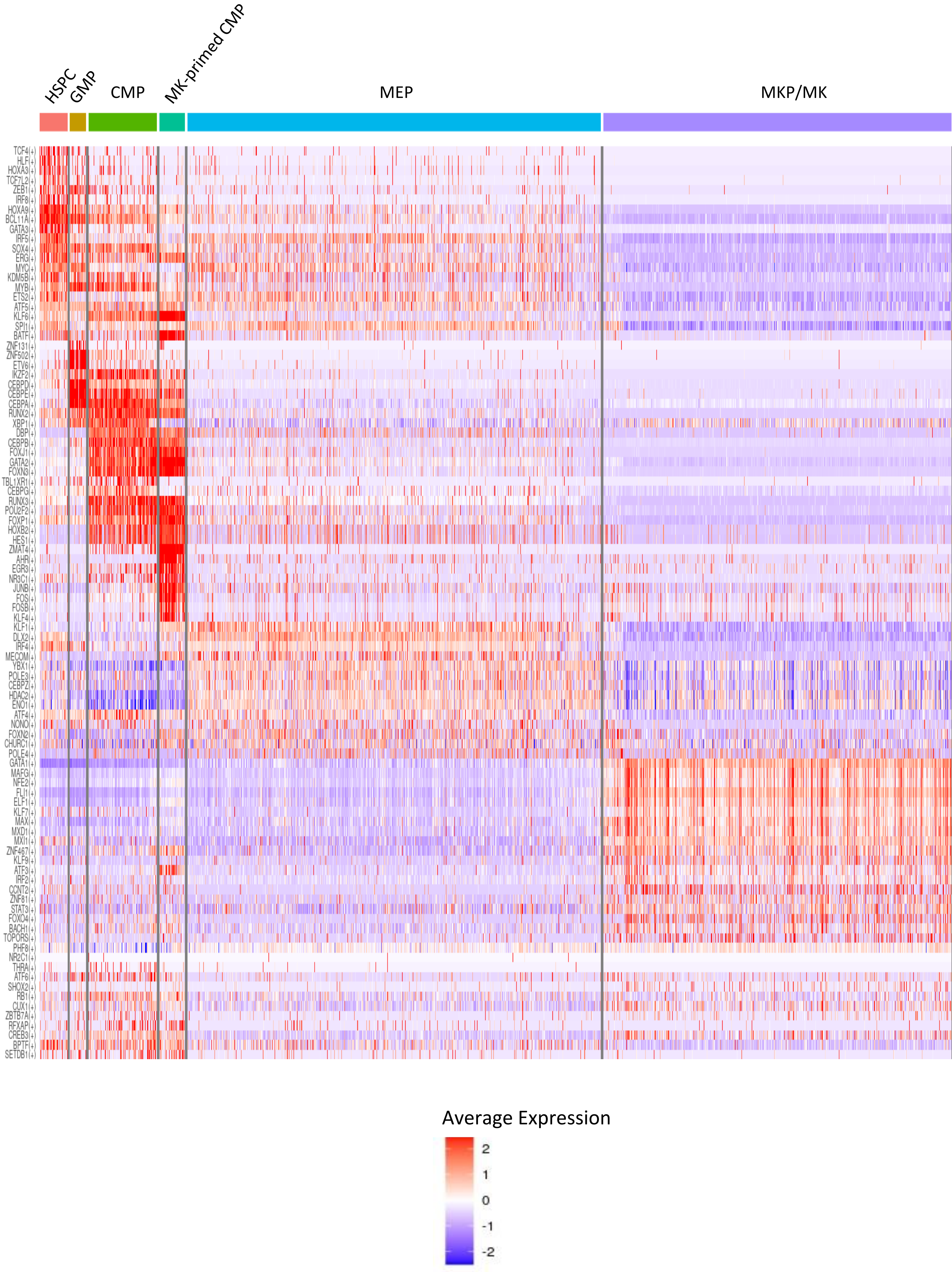

### Supplementary Figure 5

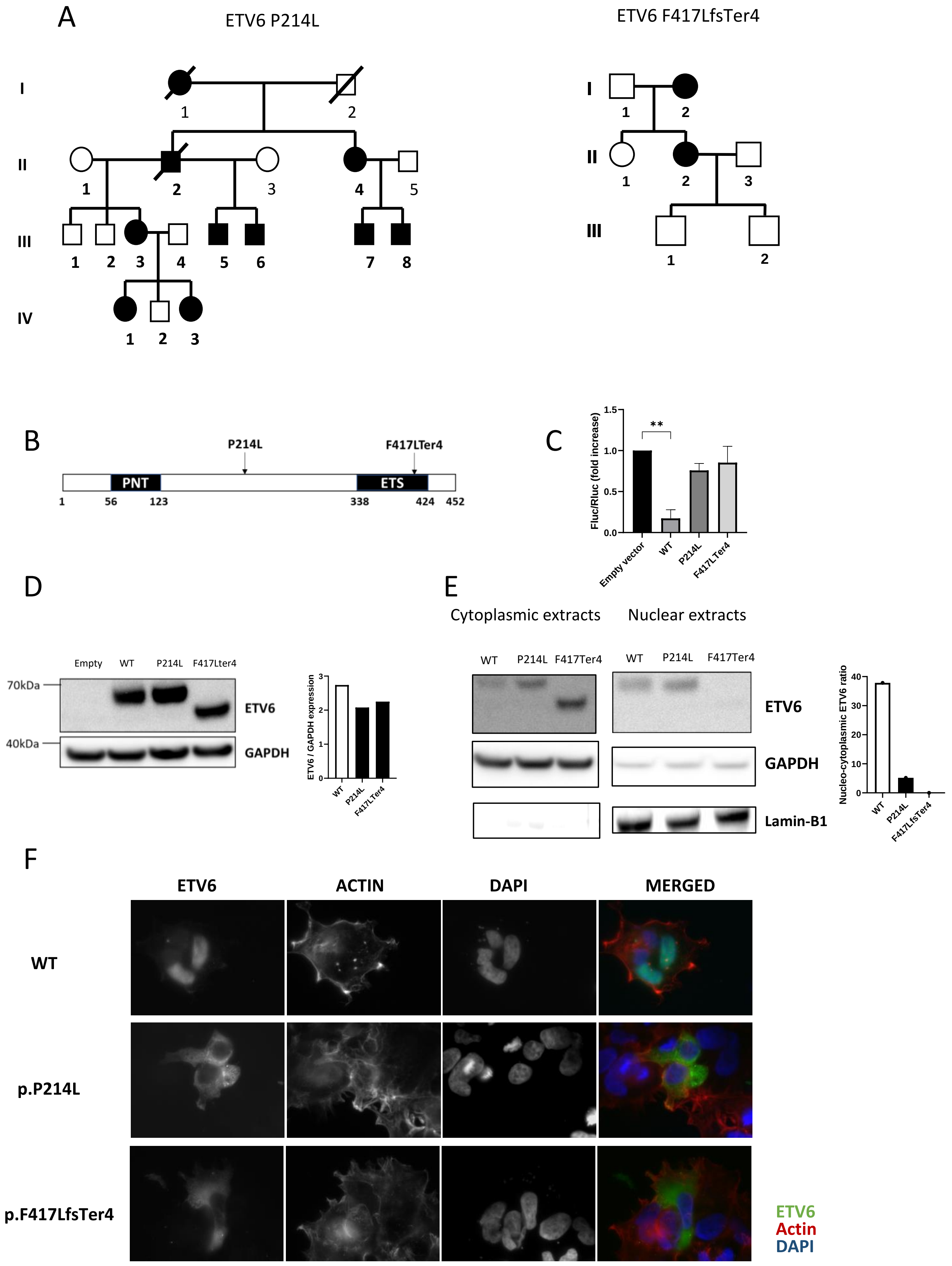

### Supplementary Figure 6

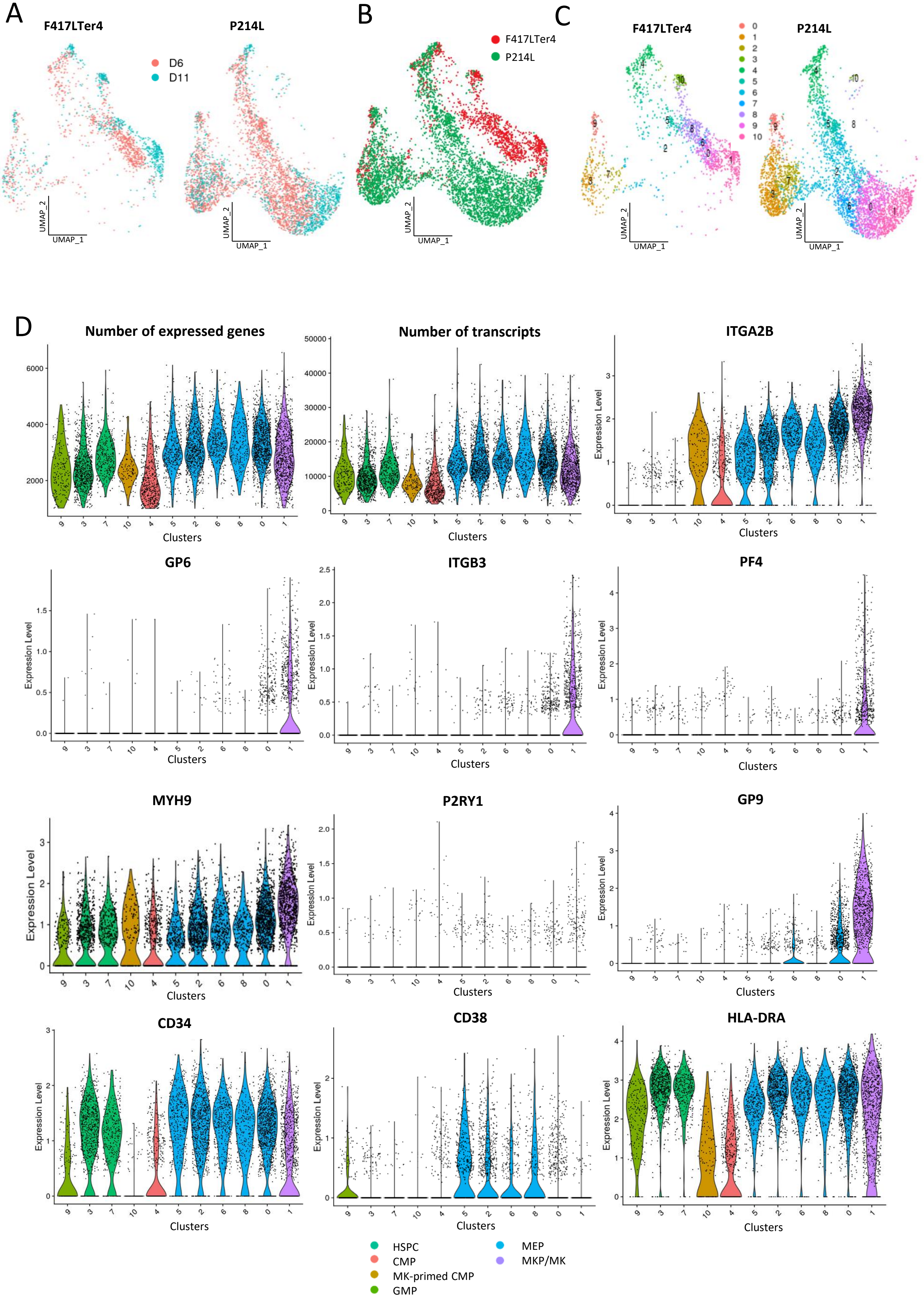

### Supplementary Figure 7

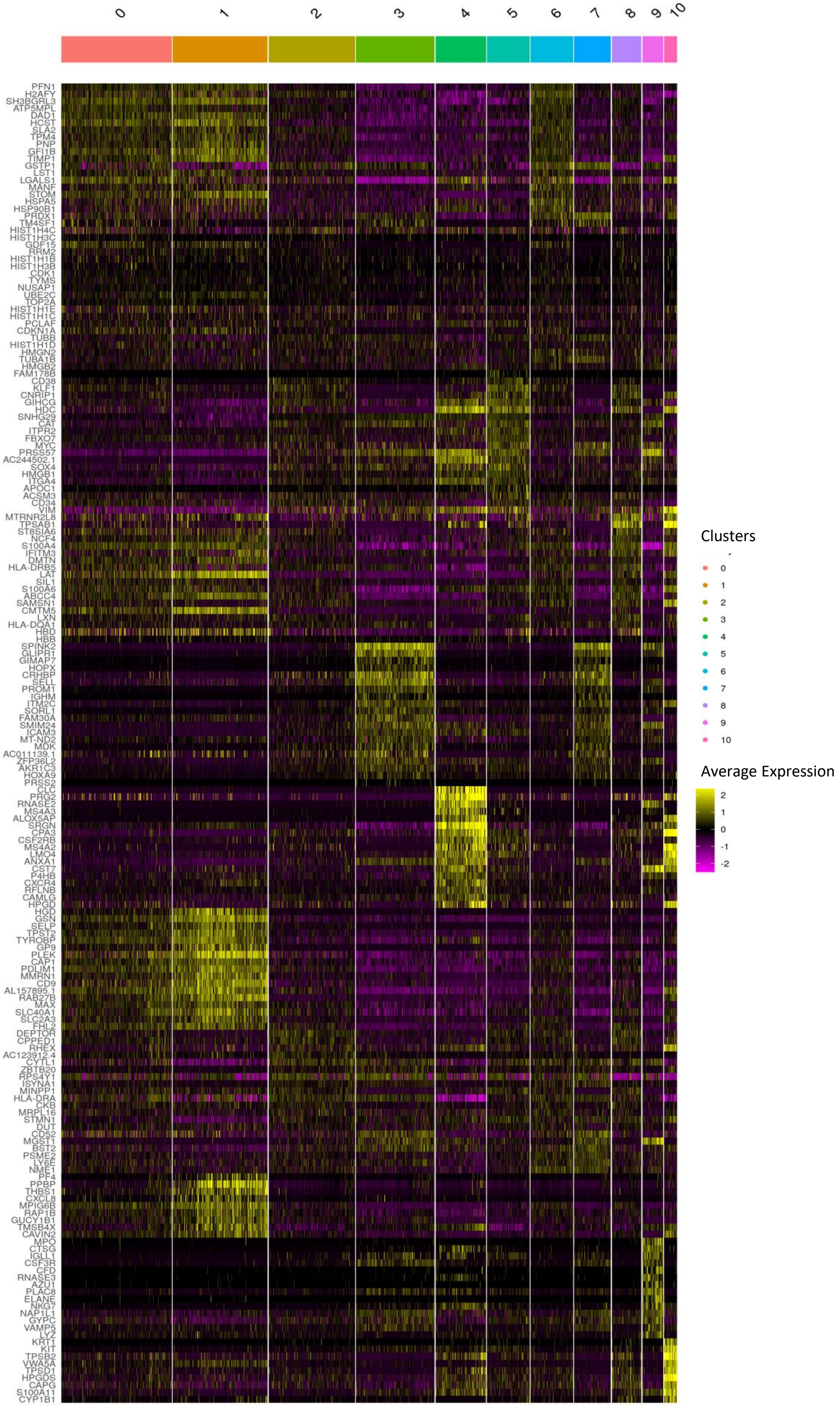

### Supplementary Figure 8

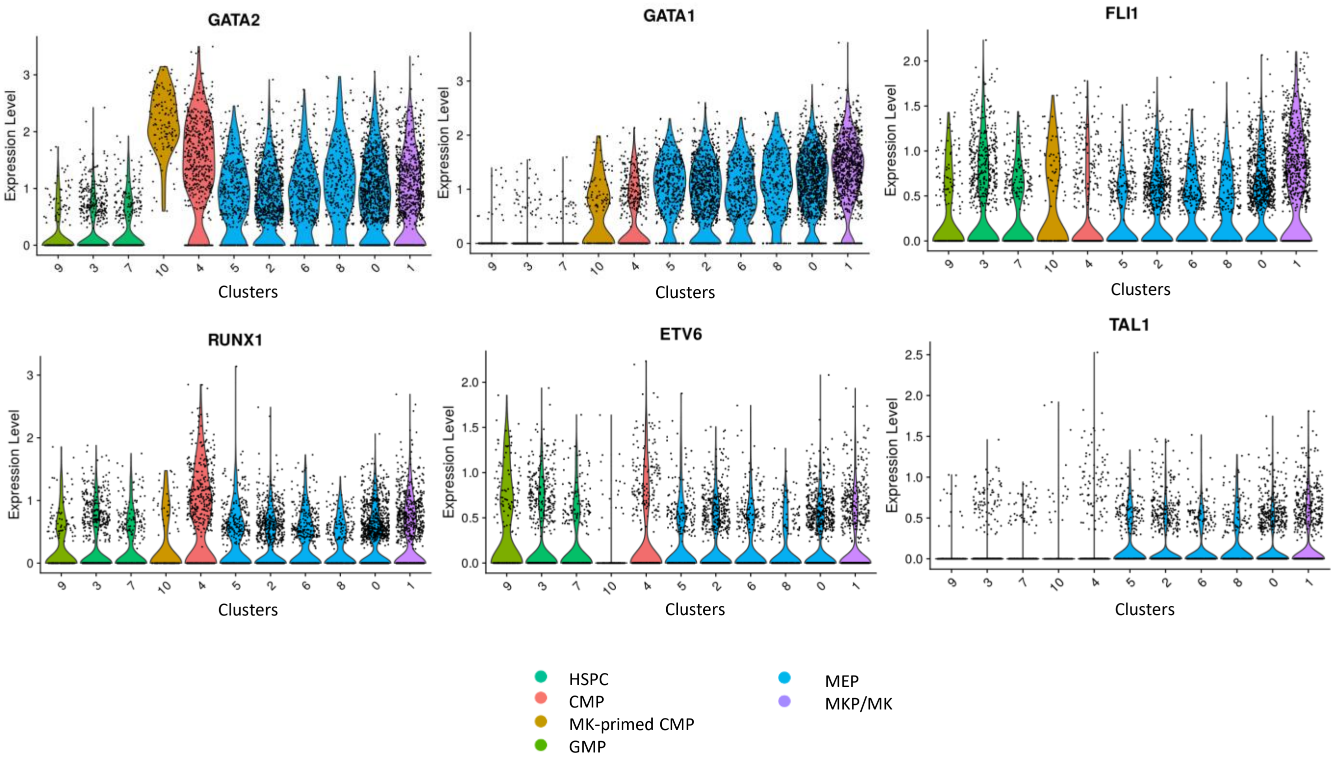

Supplementary Figure 9

A

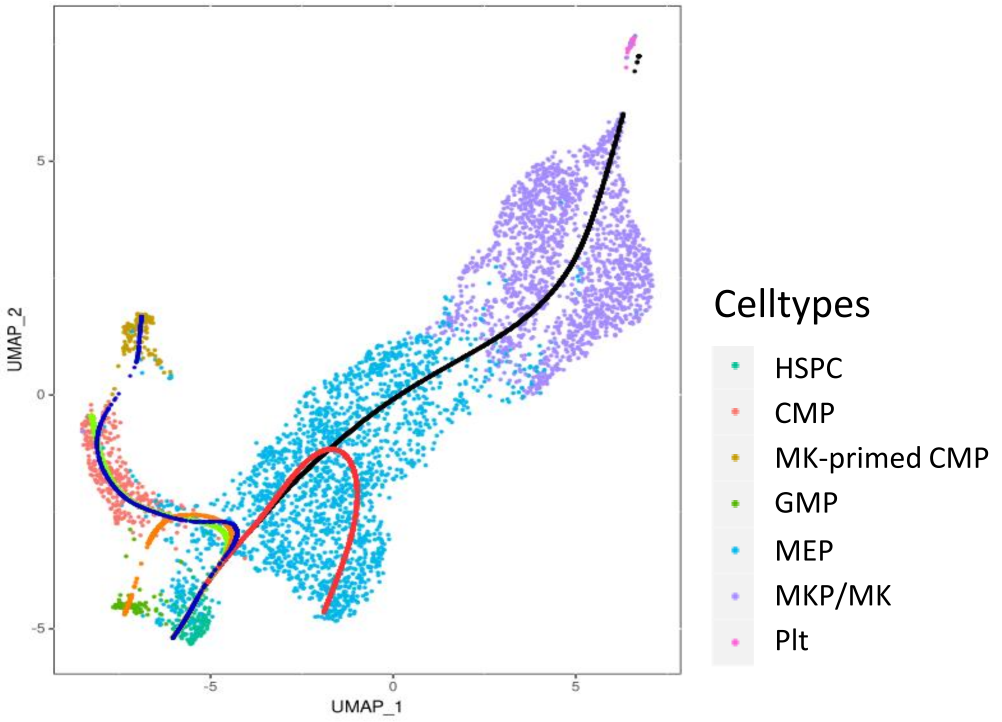


B

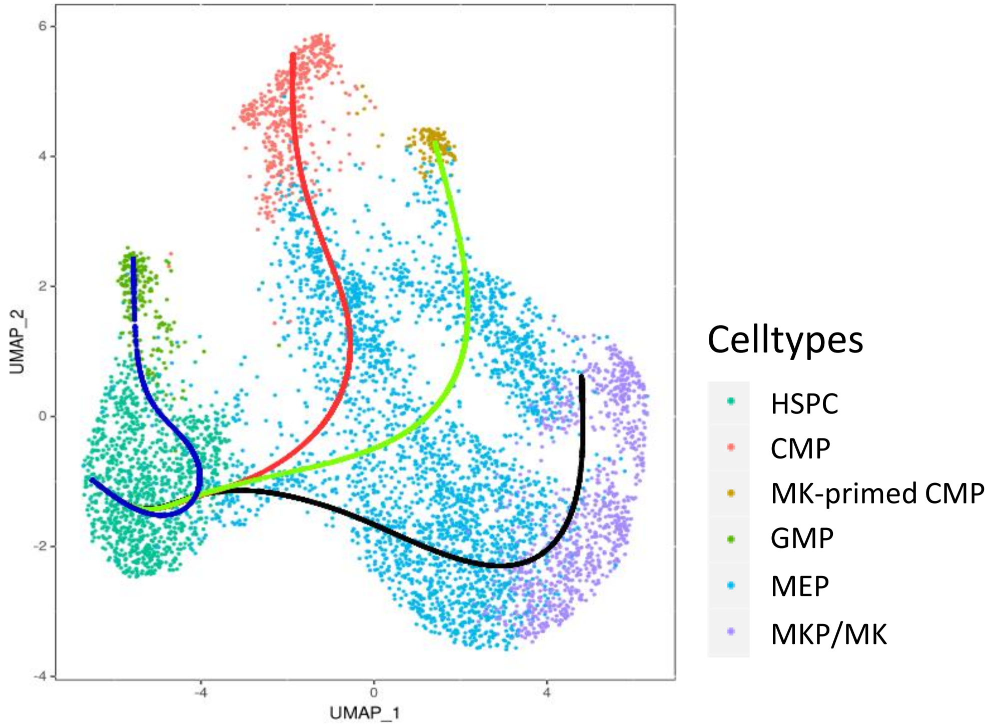

### Supplementary Figure 10

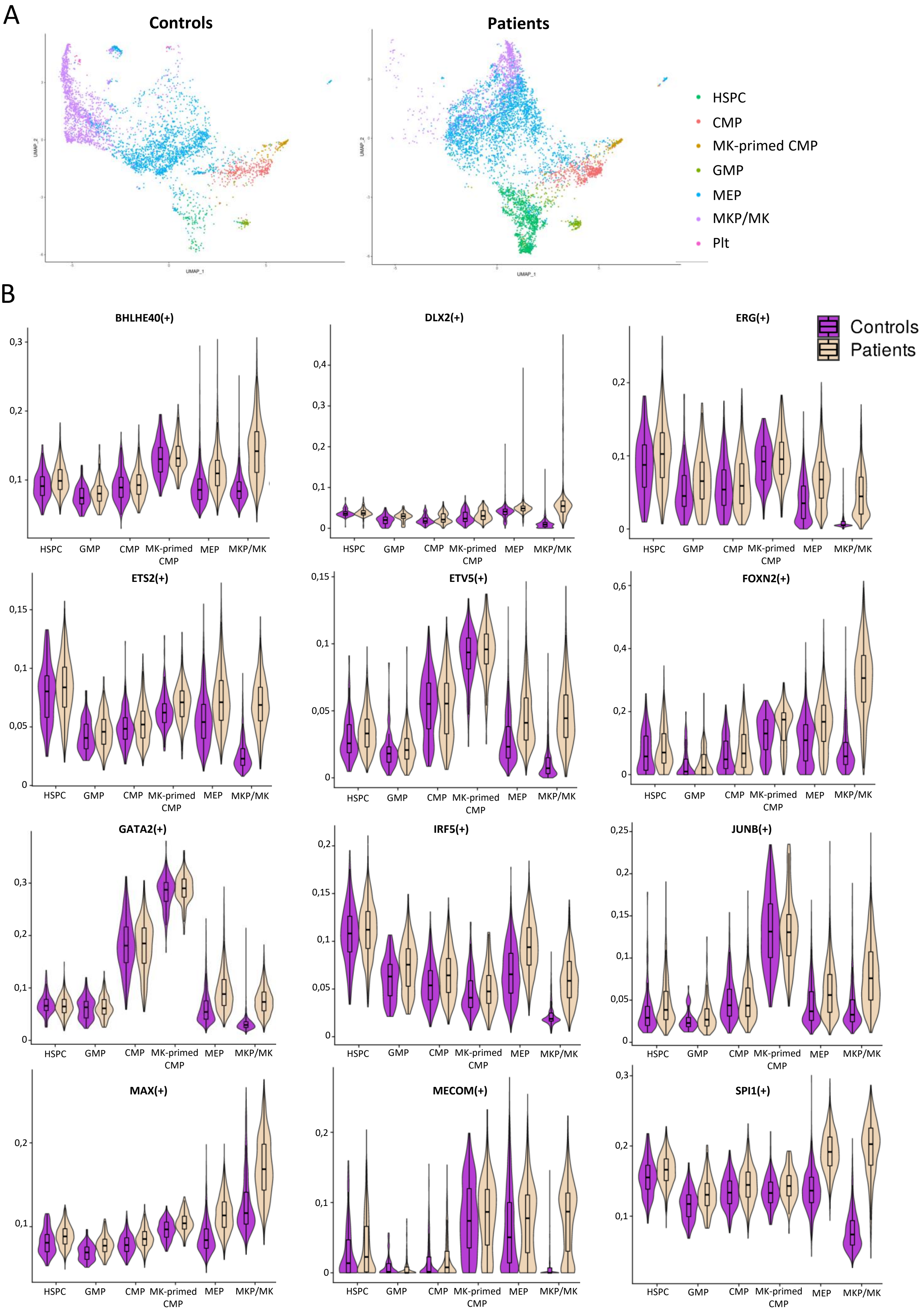

### Supplementary Figure 11

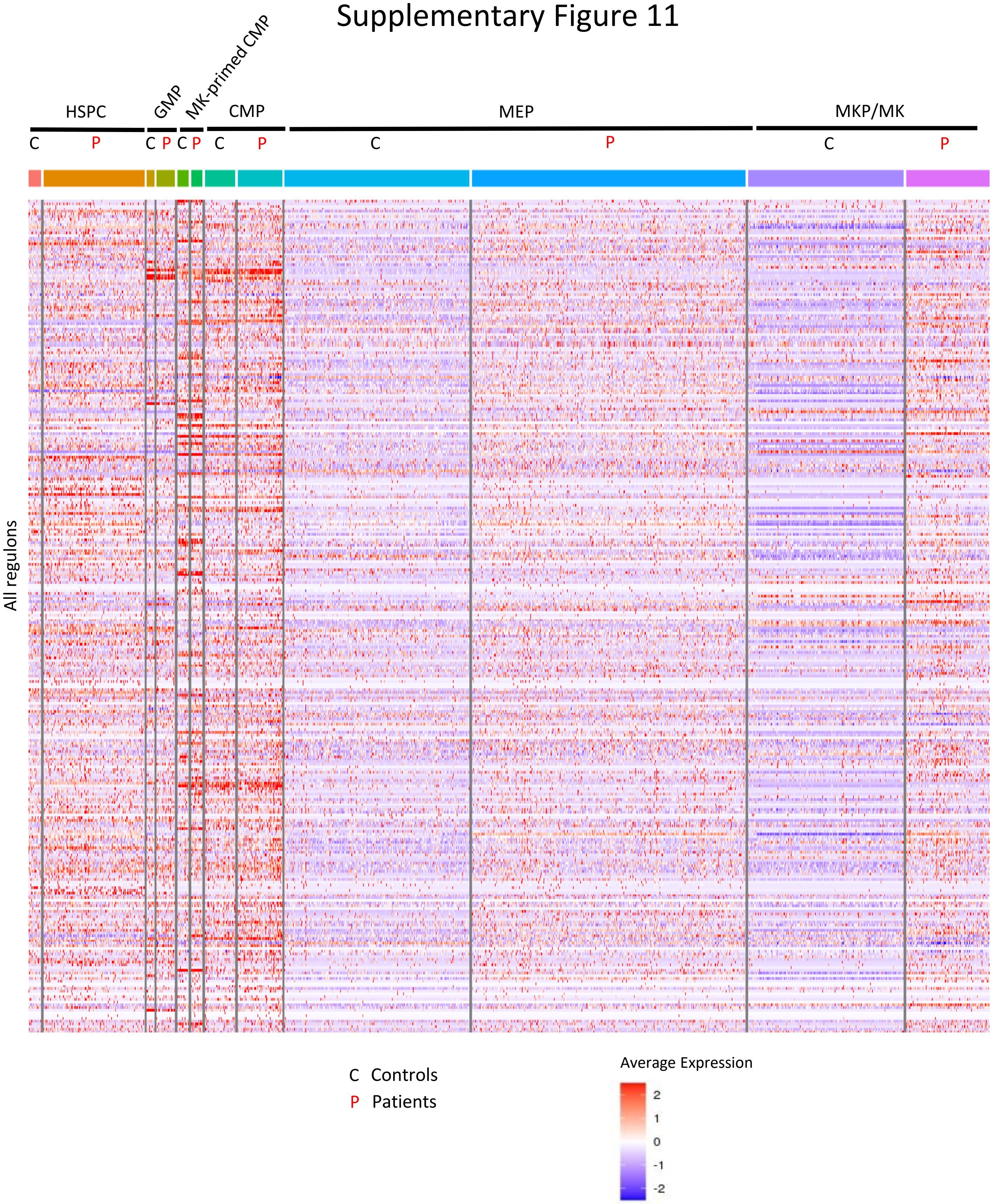

### Supplementary Figure 12

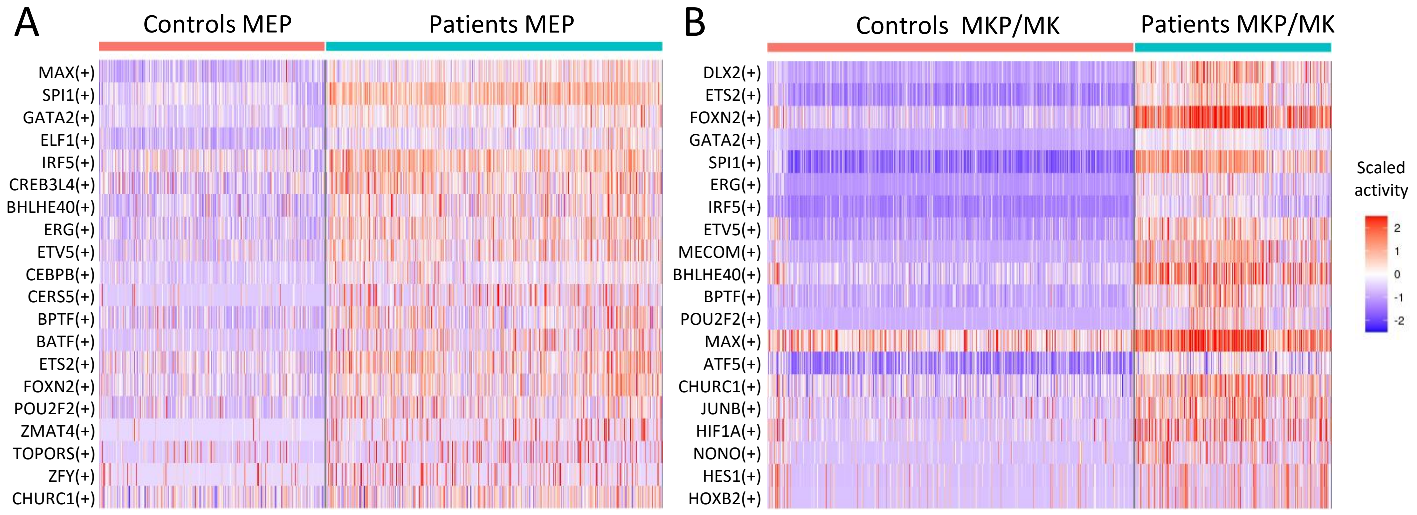

### Supplementary Figure 13

A

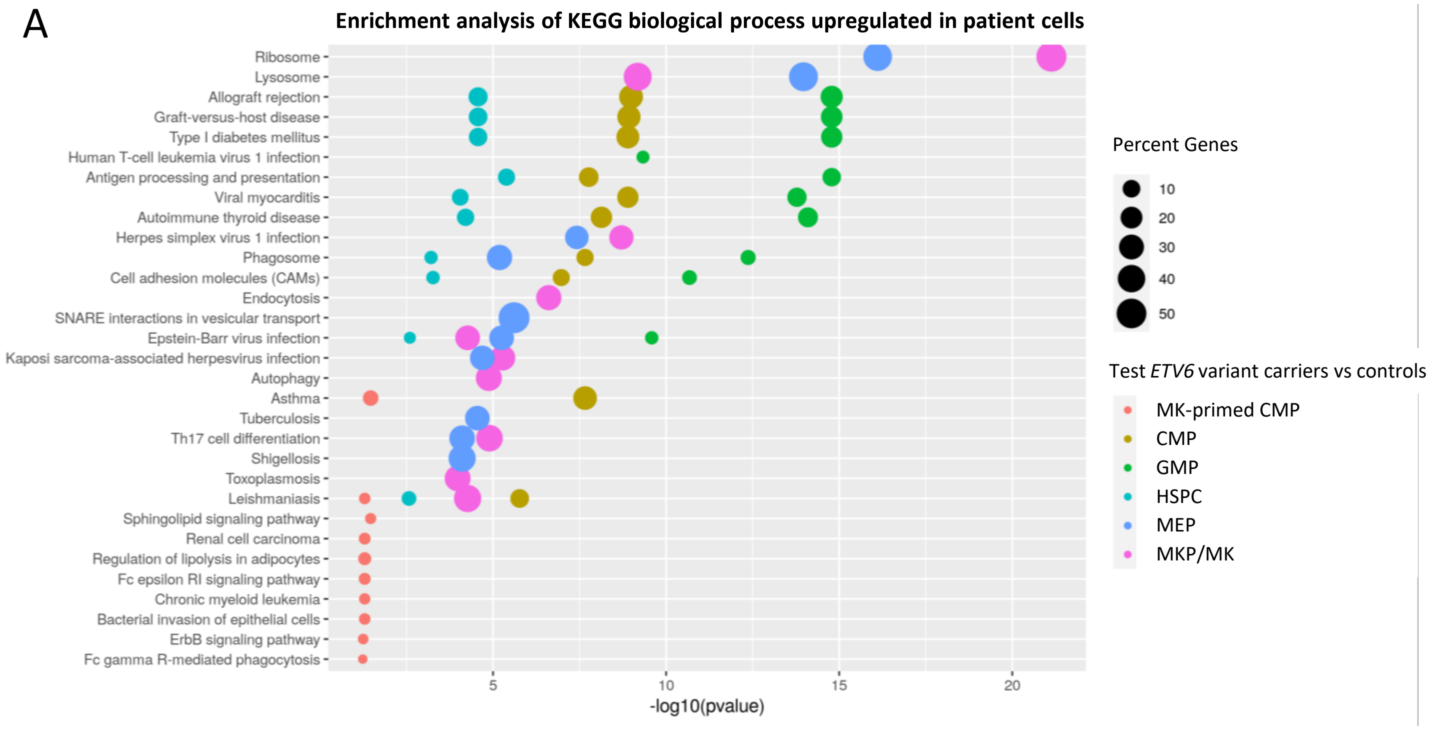


B

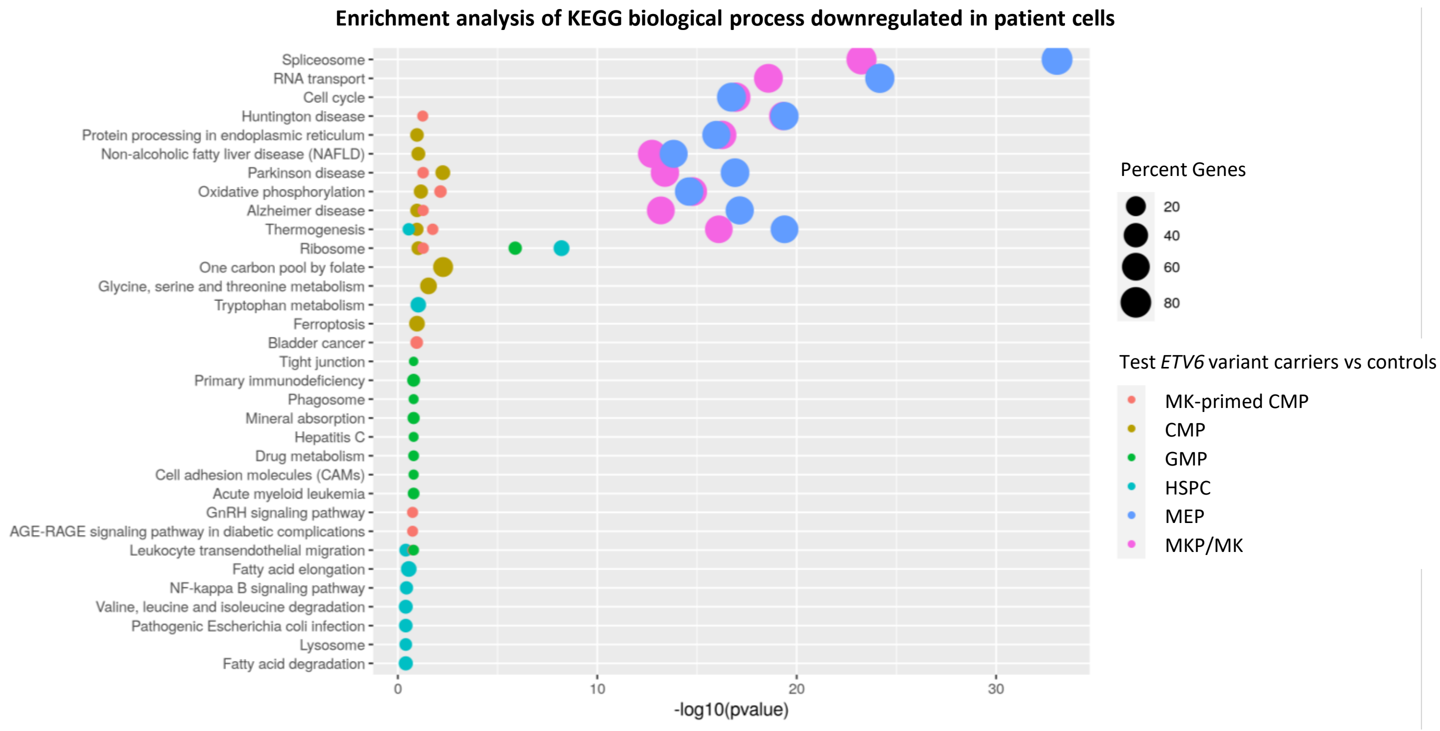

### Supplementary Figure 14

A

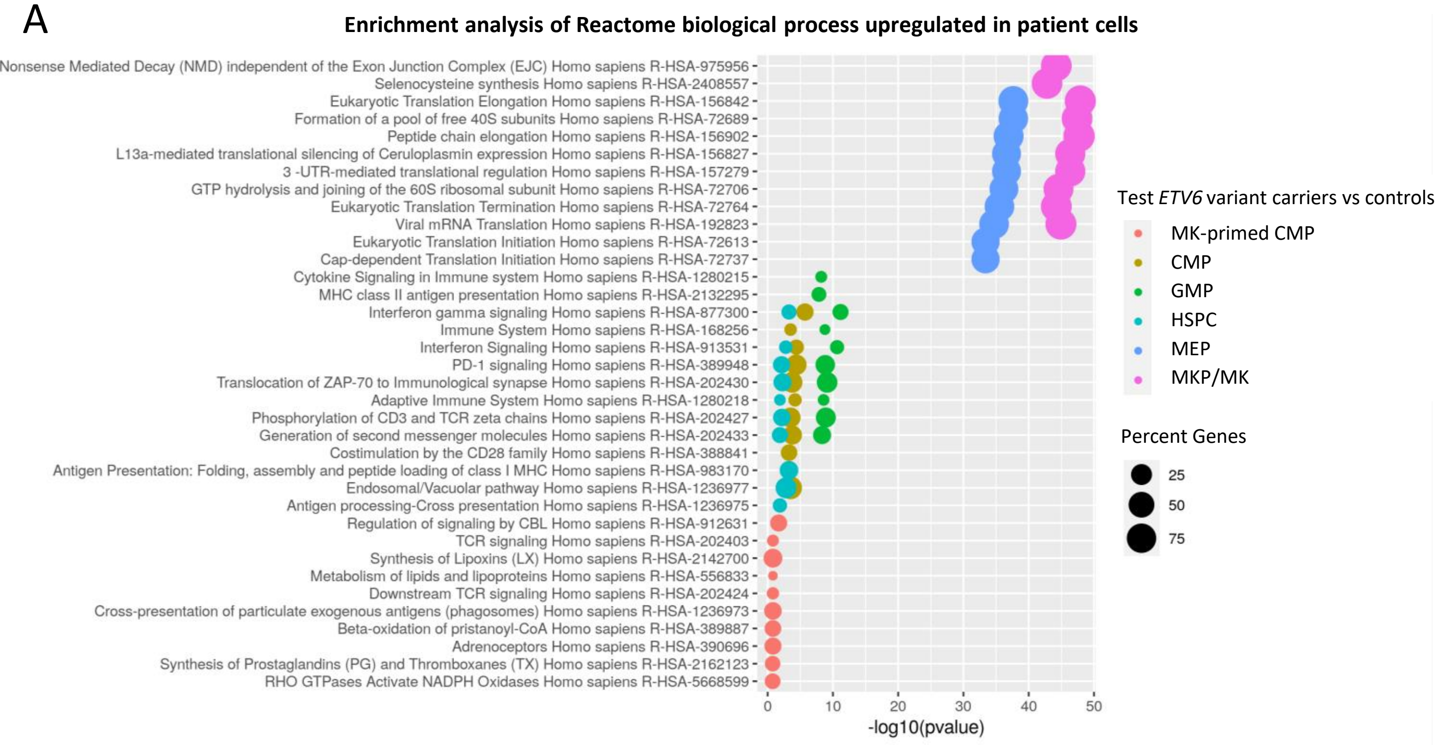


B

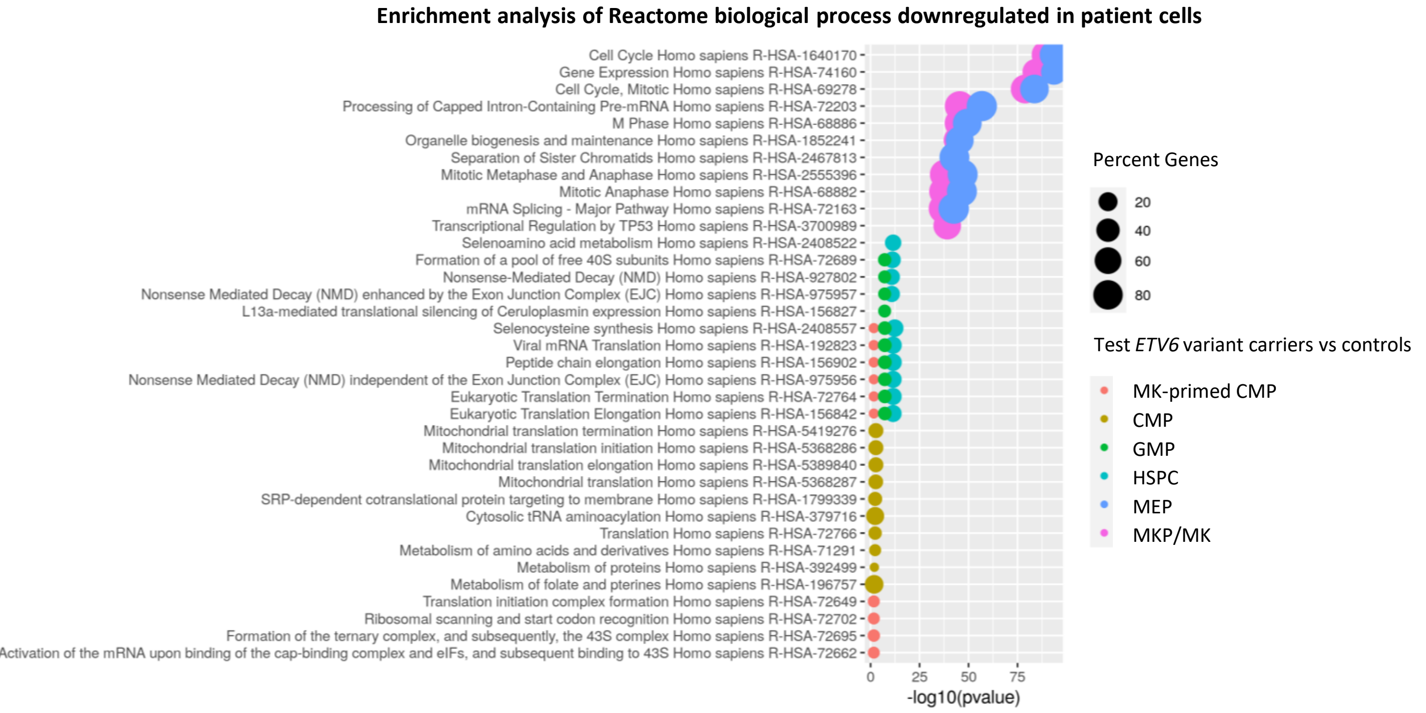

### Supplementary Figure 15

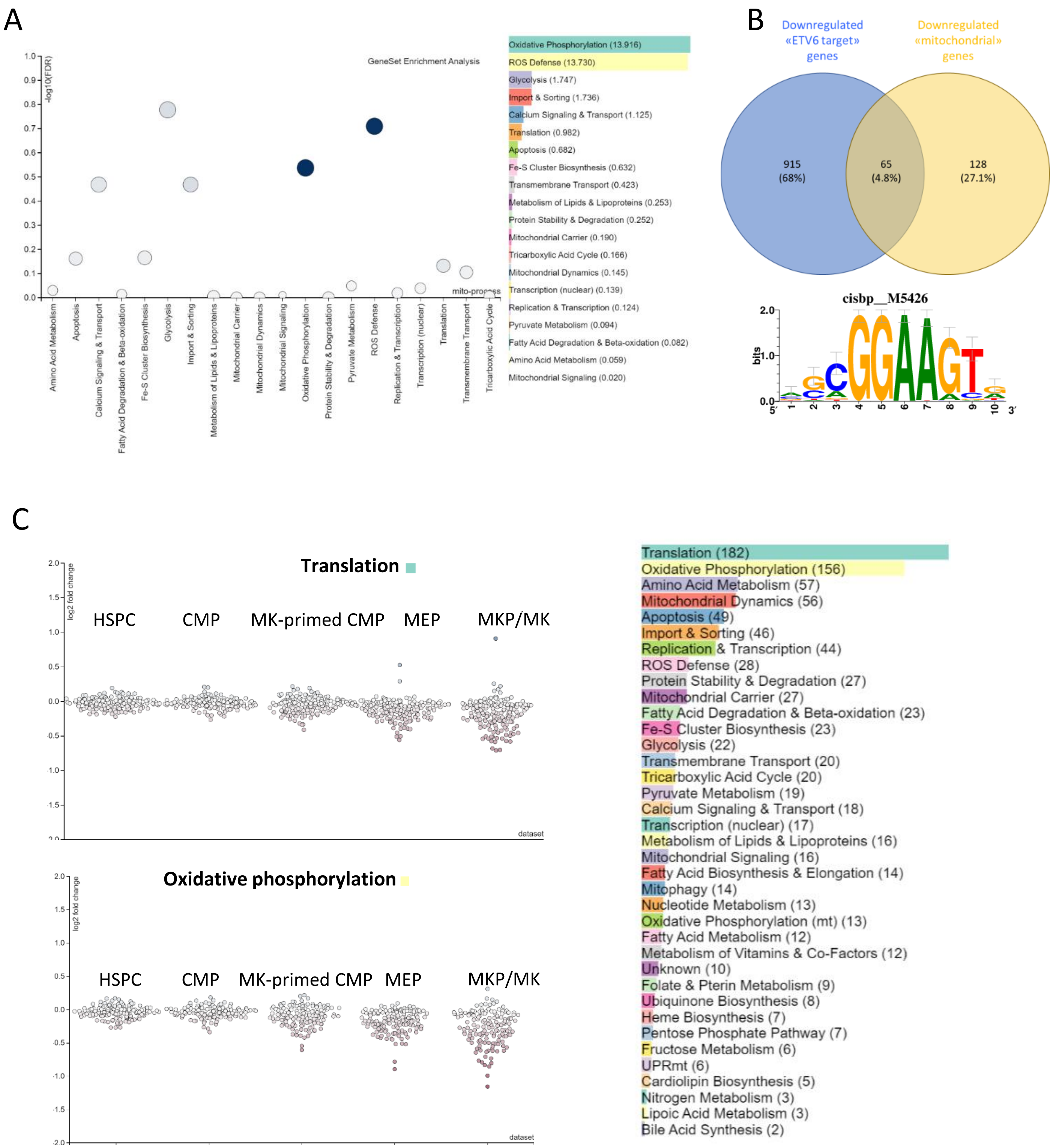

### Supplementary Figure 16

#### Translation

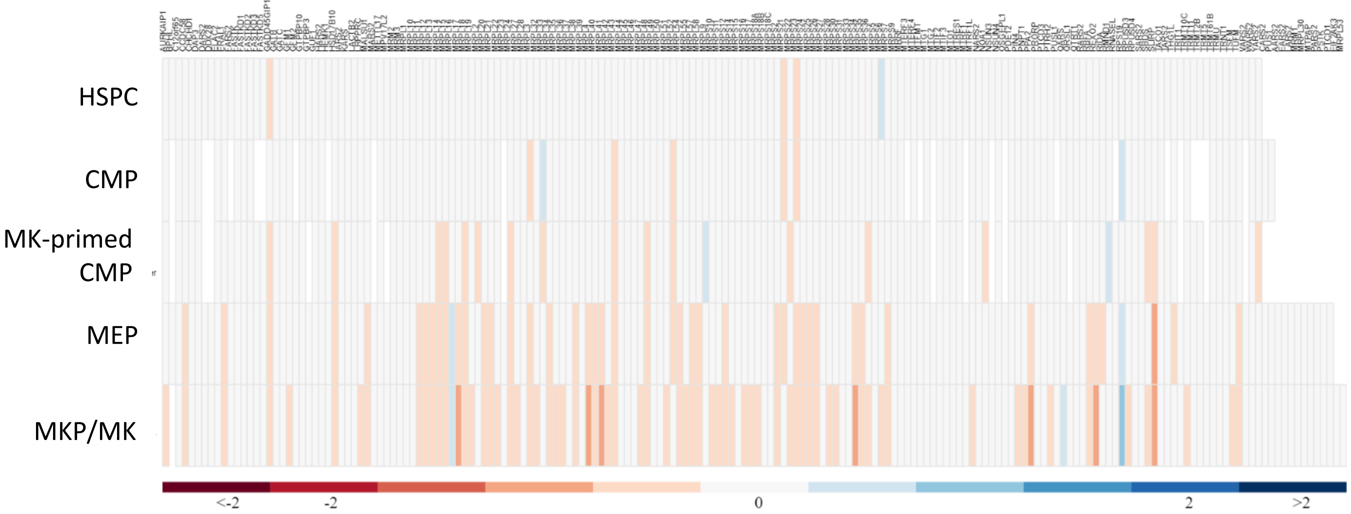


#### Oxidative phosphorylation

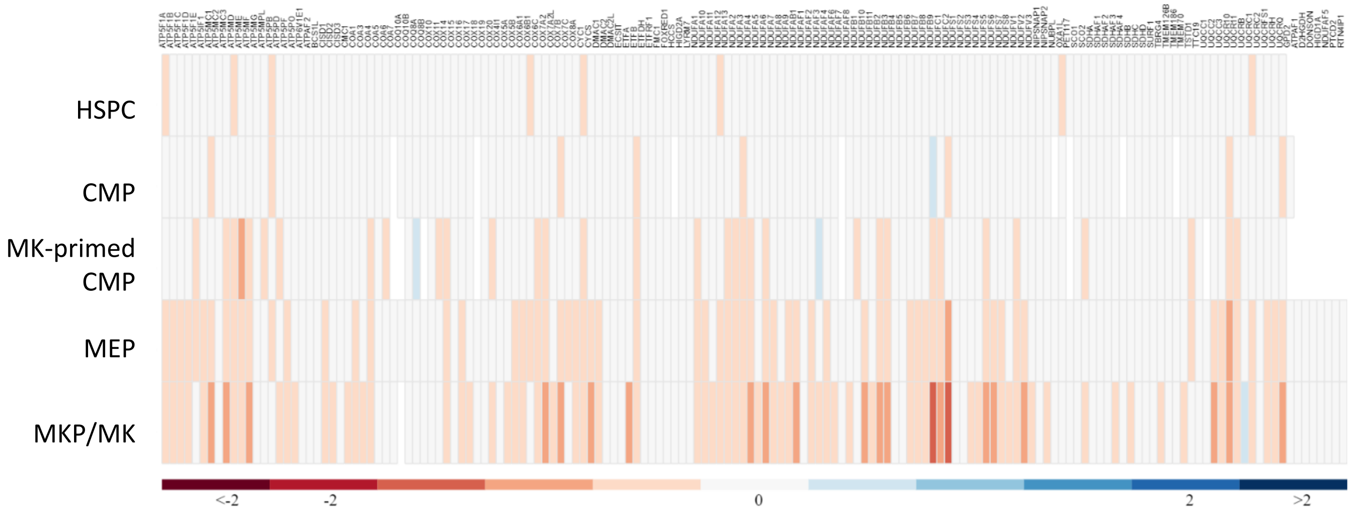

Supplementary Figure 17

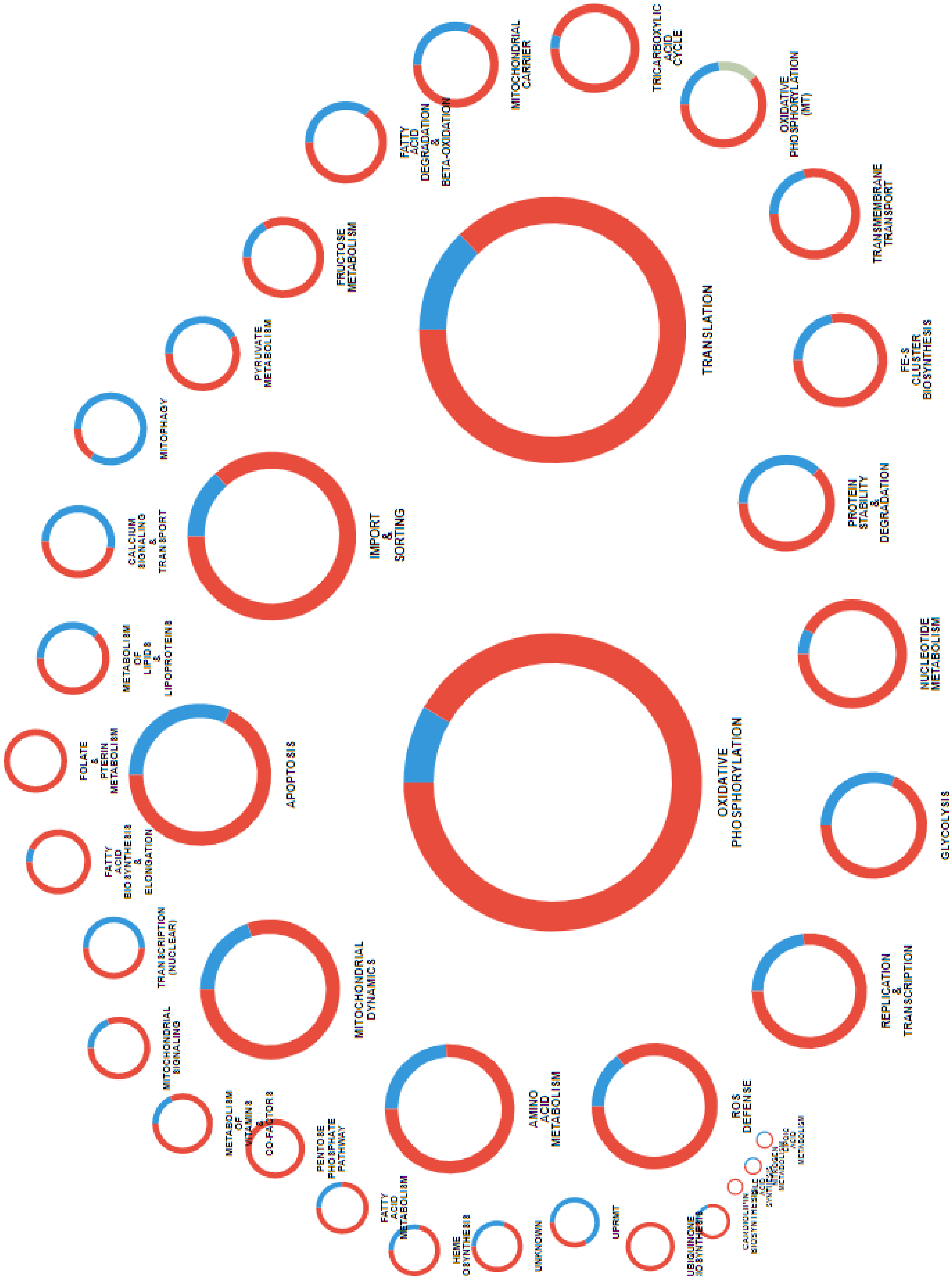

### Supplementary Figure 18

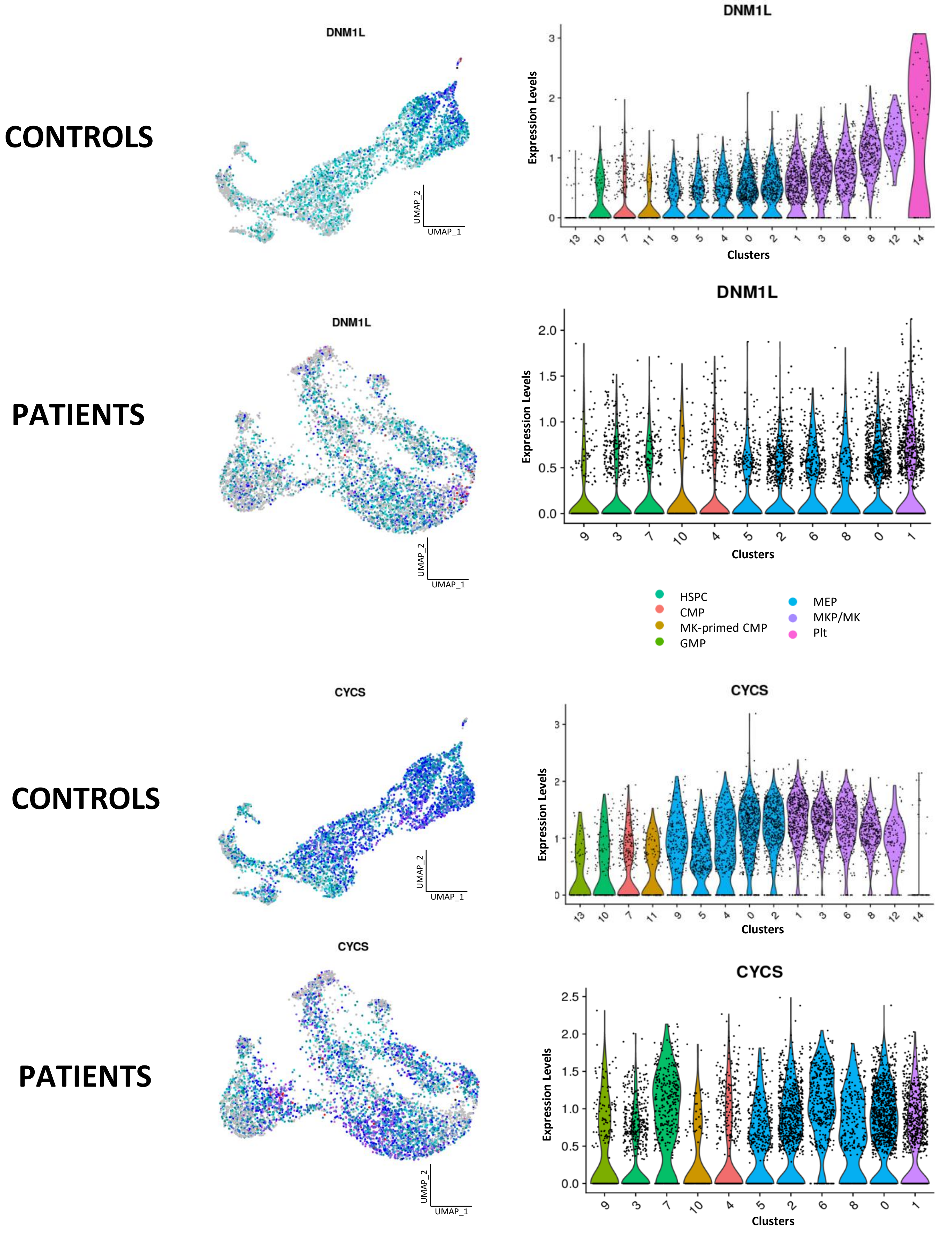

Supplementary Figure 19

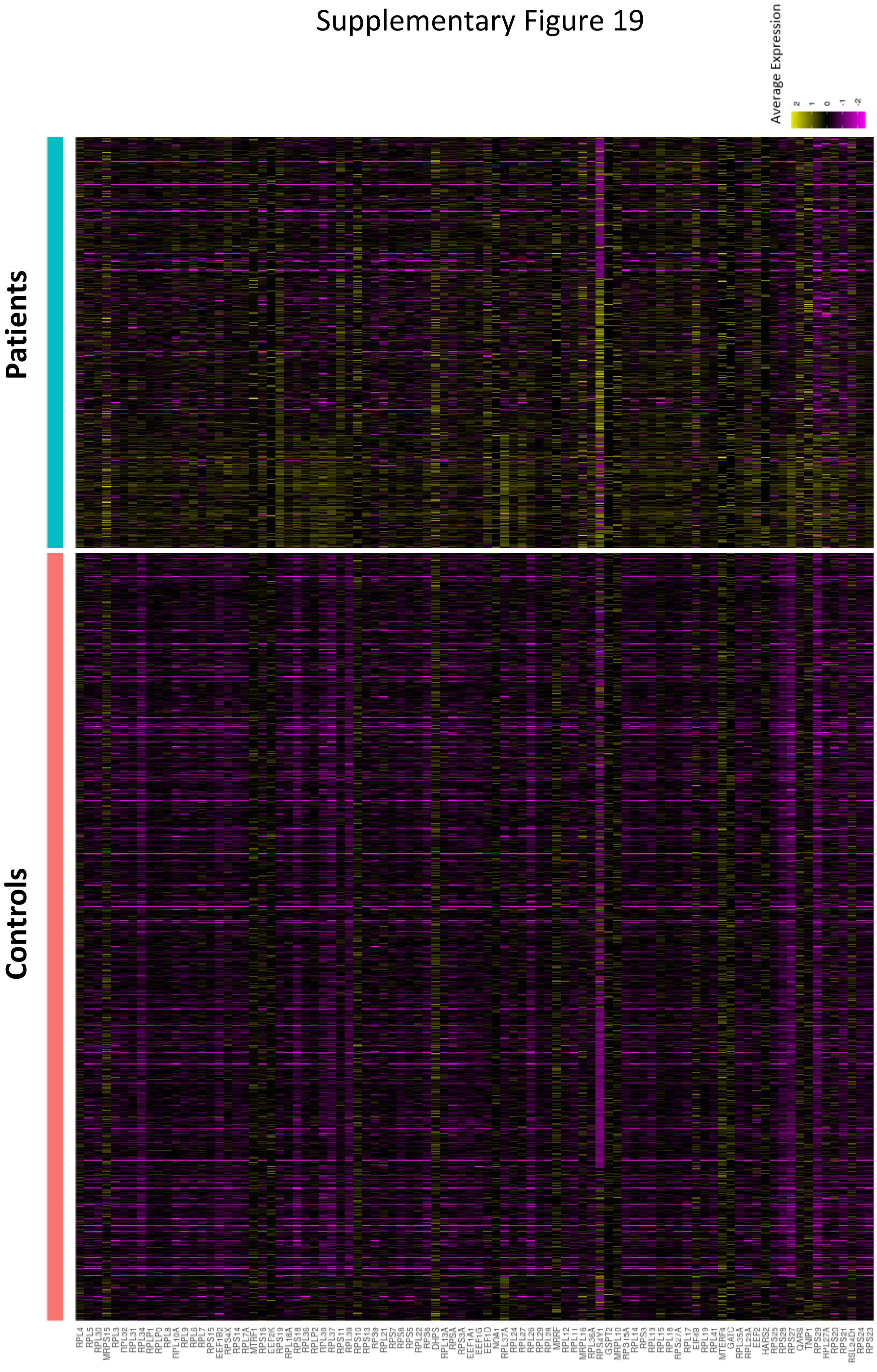

**Figures**

**Supp. figure 1: Single-cell RNA sequencing of control cells.**

A. UMAP plot of cells at days 6 and 11. Each healthy control is color-coded.

B. UMAP plot of the two demultiplexed control samples at days 6 (orange) and 11 (blue).

C. The same UMAP plot shown in Supp. figure 1B with each cluster color-coded with a distinct color.

D. Violin plots showing the number of genes expressed, transcripts and specific genes (*ITGA2B*, *GP6*, *ITGB3*, *PF4*, *MYH9*, *P2YR1*, *GP9*, *CD34*, *CD38* and *HLA-DRA*) for each of the 15 clusters. Each dot represents a single cell. Each cell type is color-coded.

**Supp. figure 2: Heatmap of the top 20 differentially expressed genes in control cells.**

Heatmap showing the average expression of the top 20 differentially expressed genes in each cluster.

**Supp. figure 3: Violin plots showing the transcription factor expression levels for each cluster (control cells).** Each cell type is color-coded.

**Supp. figure 4:** Heatmap showing top 20 regulons with differential activity between cell types in control cells.

**Supp. figure 5: Characterization of *ETV6* variants**

A. Schematic pedigrees of *ETV6*-variant inheritance. Squares represent males, circles represent females, and slashes represent deceased family members. Solid black symbols represent family members with thrombocytopenia. The F1 family carried the *ETV6* p.P214L variant, while the F2 family carried the p.F417LTer4.

B. Schematic diagram of the ETV6 protein structure. The functional N-terminal pointed domain (PNT) and C-terminal ETS DNA-binding domain (ETS) are depicted. The positions of the *ETV6* mutations are indicated.

C. GripTite 293 MSR cells were co-transfected with an empty vector, wild type (WT) or *ETV6* variant construct with the luciferase reporter plasmid containing three tandem copies of the ETS binding site upstream of the HSV TK promoter (E743tk80Luc) and the pGL4.73 Renilla luciferase control vector. Firefly:Renilla luminescence ratios (Fluc/Rluc) were calculated to assess transfection efficiency and are expressed as fold change relative to the empty vector. The data represent the mean ± SEM of four independent experiments; **P-value <0.01 vs. empty vector (one-way ANOVA with Kruskal-Wallis post-hoc test).

D. Representative western blot analysis of ETV6 expression in GripTite 293 MSR cells transfected with an empty vector, WT or *ETV6* variant constructs using a goat anti-ETV6 antibody. GAPDH was used as a protein loading control. Quantification of band intensity is shown on the right.
E. Western blot analysis of WT and ETV6 variant subcellular localization. GripTite 293 MSR cells were transfected with an empty vector, WT or *ETV6* variant constructs. Lamin-B1 and GAPDH expression levels were assessed as nuclear and cytoplasmic markers, respectively. The ETV6 nuclear/cytoplasmic ratio is shown on the right.
F. Representative immunofluorescence microscopy images of GripTite 293 MSR cells transfected with WT or *ETV6*-variant constructs visualized using bright field illumination and immunofluorescence after ETV6, actin and DAPI staining.

**Supp. Figure 6: Single-cell RNA sequencing of *ETV6*-variant cells**

A. UMAP plot of the two demultiplexed F417Lter4 and P214L patients at day 6 (orange) and day 11 (blue).
B. UMAP plot of *ETV6*-variant cells at days 6 and 11. Cells isolated from the F417Lter4 patient are shown in red, while cells from the P214L patient are shown in green.
C. The same UMAP plot shown in Supp. Figure 6B with each cluster color-coded with a distinct color.
D. Violin plots showing the number of genes expressed, transcripts, and specific genes (*ITGA2B*, *GP6*, *ITGB3*, *PF4*, *MYH9*, *P2YR1*, *GP9*, *CD34*, *CD38* and *HLA-DRA*) in each of the 11 clusters. Each dot represents a single cell. Each cell type is color-coded.

**Supp. Figure 7: Heatmap of the top 20 differentially expressed genes in *ETV6*-variant cells.**

The average expression levels of the top 20 differentially expressed genes in each cluster.

**Supp. figure 8: Violin plots showing transcription factor expression levels in *ETV6*-variant cells in each of the 11 clusters.** Each dot represents a single cell. Each cell type is color-coded.

**Supp. figure 9: Differentiation trajectories in controls and patients.**

A. UMAP plot of control cells colored by cell types. The different lines represent computed trajectories (Black = first trajectory, red = second, blue = third, green = fourth, orange = fifth).
B. UMAP plot of *ETV6*-variant cells colored by cell types. The different lines represent computed trajectories (Black = first trajectory, red = second, green = third, blue = fourth).

**Supp. Figure 10 Analysis of regulon activities display differences in MEP and MKP/MP between controls and ETV6 patients.**

A. UMAP plot superimposed with enrichment scores from the SCENIC analysis. Cells are color-coded by cell type. Lineage signature gene clustering is similar to regulon-based clustering, highlighting the same cell type-specific clusters.
B. Violin plot comparing AUC scores for each regulon across cell types. Patients are in beige and controls in purple.

**Supp. figure 11: Heatmap showing differential regulon activity between cell types derived from controls and *ETV6* patients**.

n = 312 identified regulons.

**Supp. Figure 12: The top 20 differentially active regulons.**

A. Heatmap showing the top 20 differentially expressed regulons in MEPs between controls and *ETV6* patients.
B. Heatmap showing the top 20 differentially expressed regulons in MKPs/MKs between controls and *ETV6* patients.

**Supp. figure 13: Bubble plots of the top 10 differentially expressed genes from KEGG pathways in each cell type.**

The upregulated (upper panel) and downregulated pathways (lower panel) are indicated. The cell types are color-coded.

**Supp. Figure 14: Bubble plots of the top 10 differentially expressed genes from Reactome pathways in each cell type.**

The upregulated (upper panel) and downregulated pathways (lower panel) are indicated. The cell types are color-coded.

**Supp. figure 15: Mitochondrial pathway.**

A- Using MitoXplorer 2.0, mitochondrial process enrichment analysis of *ETV6* MKPs/MKs revealed an enrichment (high combined score) in oxidative phosphorylation and ROS defense (mitochondrial processes with a high combined score are highlighted in dark blue).

B- Venn diagram of downregulated genes with an ETV6 binding site (blue circle) and mitochondrial pathway genes (yellow circle) in MKPs/MKs isolated from controls vs. *ETV6* patients. Among all downregulated genes harboring an ETV6 binding site, 65 genes are associated with mitochondrial pathways.

C- Scatter plots, a sortable heatmap and a mitochondrial processes bar chart are displayed. The first two deregulated mitochondrial processes are displayed: translation (upper panel) and oxidative phosphorylation (lower panel). The scatter plot shows the log2 fold change (y-axis) and datasets (x-axis). Each bubble represents one gene, in which downregulated genes (red bubbles) and upregulated genes (blue bubbles) are indicated.

**Supp. Figure 16: Heatmap showing the individual genes and datasets for translation and oxidative phosphorylation pathways (MitoXplorer).**

The genes are color-coded according to log2 fold change and classified by cell types.

**Supp. Figure 17: MitoXplorer results: interactome view.**

Overview of all mitochondrial processes of the MKP/MK dataset. Each process is shown as a circle with colored segments according to the proportion of dysregulated genes. Upregulated genes are shown in blue, and downregulated genes are shown in red.

**Supp. Figure 18: Downregulated mitochondrial genes *DNML1* and *CYCS* in *ETV6*-variant cells**

UMAP plots and violin plots showing the expression of the *DNML1* and *CYCS* genes in all clusters. Patient and control cells are shown.

**Supp. Figure 19: Heatmap of the differentially expressed genes associated with "translation pathways" in *ETV6*-variant megakaryocytes**

The most striking upregulated translation pathway genes included primarily *RPS* and *RPL*, which encode ribosomal protein small and ribosomal protein large subunits, respectively.
